# Phenotypic heterogeneity and kidney tropism of *Klebsiella pneumoniae* clinical urinary tract infection isolates

**DOI:** 10.64898/2026.03.27.712997

**Authors:** Grace E. Shepard, Zachary Mills, Drew A. Pariseau, Brooke E. Ryan, Jolie Lagger, Laura A. Mike

## Abstract

Urinary tract infections (UTIs) are a significant public health burden that impact millions of people every year and are highly prevalent among in hospital-acquired infections. *Klebsiella pneumoniae* is the second most common cause of UTIs after uropathogenic *Escherichia coli* (UPEC). Thus far, the molecular mechanisms underlying pathogenesis is better understood in UPEC than *K. pneumoniae*. UPEC is known to have fitness factors such as fimbrial adhesion and evasion of complement-mediated killing. In other infection types, *K. pneumoniae* fitness has been associated with mucoidy and diverse capsular serotypes. To establish *K. pneumoniae* virulence factors contributing to UTI, we examined how environmental cues regulate urovirulence-associated phenotypes in clinical *K. pneumoniae* UTI strains. These factors included capsular polysaccharide properties, hemagglutination, serum resistance, adherence to bladder epithelial cells, and *in vivo* fitness. We found that clinical *K. pneumoniae* UTI isolates phenotypes are highly heterogeneous and can change in response to human urine. Despite *K. pneumoniae* clinical isolates presenting heterogeneous fitness properties, all similarly colonize the urinary tract. These results suggest that additional fitness factors contribute to *K. pneumoniae* uropathogenesis. Identifying these shared fitness factors will provide mechanistic insights into *Klebsiella* uropathogenesis and reveal candidate therapeutic targets.

## INTRODUCTION

Urinary tract infections (UTIs) are one of the most common bacterial infections, with over 400 million infections and over 230 thousand deaths occurring annually.^1, 2^ The impact of UTIs is particularly significant for women, with over 50% being diagnosed with a least one UTI in their lifetime.^3^ With an estimated 25-40% of primary care antibiotic courses being prescribed to treat UTIs, the rise in antibiotic resistance threatens our ability to treat these common infections.^4^ Therefore, it is pertinent to define the bacterial mechanisms used to cause UTIs.

*Klebsiella pneumoniae* frequently causes UTIs and is the second and third leading causes of uncomplicated and complicated UTIs, respectively.^5^ The World Health Organization (WHO) lists *K. pneumoniae* as the top bacterial threat on the WHO Bacterial Priority Pathogens List, partially due to its accumulation of multi-drug resistance genetic determinants.^6^ Despite causing a variety of infections, *K. pneumoniae* is most frequently isolated in hospital settings from urine cultures.^7^ There are two *K. pneumoniae* pathotypes: classical (cKp) and hypervirulent (hvKp). cKp are nosocomial strains that primarily infect immunocompromised patients and are more prevalent in UTIs, whereas hvKp are community-acquired strains that can infect immunocompetent patients and often lead to more severe disease.^8–10^ Even though hvKp encode more virulence factors, cKp strains are more frequently isolated from UTI patients than hvKp strains.^11^ Despite its prevalence, cKp strains are understudied in UTIs compared to the leading cause of UTIs, uropathogenic *Escherichia coli* (UPEC).^12^

Our current understanding of uropathogenesis has been primarily based on UPEC studies. These works have shown that gut colonization by uropathogenic bacteria typically precedes UTIs.^13^ If UPEC is introduced to the periurethral area, it can colonize and ascend the urethra.^13^ Once UPEC traverses the urethra, it then causes ascending infections of the bladder, kidneys and, if left uncontrolled, the bloodstream.^13^ Many virulence factors have been characterized in UPEC; including adhesins such as type 1 and type 3 fimbriae, and immune evasion factors such as capsule which blocks complement-mediated killing.^14^ Similar to UPEC, *K. pneumoniae* type 1 fimbriae are broadly conserved among diverse clinical and environmental isolates, with 90% expressing *fimH*, a type 1 fimbriae tip adhesin important for bladder epithelial cell binding.^15, 16^ However, there remains a critical knowledge gap in understanding what virulence factors cKp has compared to UPEC virulence factors and whether these phenotypes impact *in vivo* uropathogenesis.

While *K. pneumoniae* shares some virulence factors with UPEC, including capsule, *K. pneumoniae* relies more heavily on capsule for virulence and more frequently exhibits mucoidy.^17^ *K. pneumoniae* employs several capsule-dependent methods to evade the innate immune system, including complement resistance and reduced phagocytosis, which can be quantified by serum survival and macrophage association assays.^18^ Capsule serotypes K1 and K2, more often associated with hvKp strains, generally display greater virulence compared to other capsule serotypes due to enhanced immune evasion.^17^ In contrast to systemic infections, capsule serotypes among *K. pneumoniae* UTI strains are highly diverse, with one study reporting 23 different K serotypes from 32 UTI isolates from elderly patients.^19^ The mucoid phenotype is attributed to increased capsular polysaccharide (CPS) chain length uniformity and length, which increase sedimentation resistance.^20, 21^ These long, uniform CPS chains produce tacky colonies and reduce macrophage binding.^21^ As such, mucoidy is typically associated with invasive hvKp infections and increased virulence; however, UTIs are predominantly caused by non-mucoid cKp strains.^22^ Interestingly, hvKp strains suppress mucoidy in response to urine compared to growth in Luria-Bertani (LB) broth.^21^ How cKp strains respond to environmental cues encountered in urine compared to hvKp remains unknown. Thus, there remains a significant gap in understanding how *K. pneumoniae* virulence factors are regulated in the context of uropathogensis.

To examine whether cKp UTI isolates express known virulence factors, we determined if clinical cKp UTI isolates change their mucoidy and capsule abundance based on environmental cues relative to two hvKp controls. We used urine to simulate the bladder environment and LB medium for a nutrient-rich control environment. We then examined how exposure to both media impacts other urovirulence-associated phenotypes including: hemagglutination, serum resistance, and adherence to bladder epithelial cells. Ultimately, we assessed the *in vivo* fitness of four strains, which expressed varying combinations of virulence phenotypes. We found that *K. pneumoniae* clinical isolates exhibited heterogeneous fitness phenotypes that were primarily attributed to inter-strain variation and not differentially regulated between LB and urine. However, we consistently observed that urine upregulated capsule abundance, which correlated with increased serum resistance. Despite broad phenotypic heterogeneity, clinical *K. pneumoniae* strains all established robust UTI in a murine model, consistently causing pyelonephritis. This study highlights the heterogeneity of *K. pneumoniae* clinical UTI isolates and advances our understanding of *K. pneumoniae* uropathogenesis.

## RESULTS

### Genomic analyses of clinical *Klebsiella* UTI isolates

Clinical isolates were collected from patients at the University of Michigan Medical Center.^23–25^ Patients colonized by *Klebsiella* based on a rectal swab were followed longitudinally for subsequent clinical culture of *Klebsiella* isolates. Patients with positive urine cultures and symptoms meeting CDC National Healthcare Safety Network UTI case definitions were diagnosed with an active UTI.^23, 26^ Twenty-five *Klebsiella* isolates that met the clinical definitions for UTI were selected for genomic and phenotypic analyses, and served as cKp strains (**Tables 1 and S1**). Two model hvKp strains, KPPR1 and NTUH-K2044, were included as points of comparison to the classical strains selected for this study.

Genomic analyses of the UTI isolates using Pathogenwatch revealed that 21 UTI isolates were *K. pneumoniae subsp. pneumoniae*, three isolates (Kp6486, Kp10079, and Kp10081) were *K. pneumoniae subsp. variicola*, and one isolate (Kp10403) was *K. pneumoniae subsp. quasipneumoniae* (**Table 1**).^27–29^ According to Pathogenwatch, all *K. pneumoniae* UTI isolates had low virulence scores (0-1) and did not encode the *rmp* locus (**Table 1**). The *rmp* locus is often encoded by hvKp strains and is responsible for increased capsule production and mucoidy.^30^ Homologs of known adhesins were identified in the *Klebsiella* UTI isolates by performing BLAST analyses using adhesin sequences from UPEC strains.^31^ All isolates, except Kp7951, encoded FimA homologs with 81-83% identity, PapA homologs with 25–31% identity, SfaA homologs with 57–65% identity, and MrkA homologs with 93-95% identity to UPEC adhesin sequences (**Tables 1 and S2**). Kp7951 encodes a FimA homolog with 30–32% identity, no PapA homolog, a SfaA homolog with 35% identity, and a MrkA homolog with 26% identity to UPEC. Overall, BLAST analysis revealed that cKp isolates had high adhesin homology with UPEC.

**Table 1.** Genomic characterization of clinical *Klebsiella* UTI Isolates. Genomes were assembled in PATRIC then uploaded to Pathogen watch for species identification and characterization. Isolates with ≥25% identity to adhesin sequences present in UPEC are listed.

| Strain | Species | Sequence Type | K Locus | Predicted Capsule Type | O Locus | Predicted O Type | Antibiotic Resistances | Virulence Score | Hypermucoidy (RmpADC/*rmp* A2) | Plasmid Types | Fimbriae/Adhesins (ECC/ECI/MS2027) |
| --- | --- | --- | --- | --- | --- | --- | --- | --- | --- | --- | --- |
| KPPRI | *K. pneumoniae* | 493 | KL2 | K2 | O1/O2v1 | O1 | Penicillin (SHV-1 (homolog)) | 1 - yersiniabactin only | rmp 3; ICEKp1 / - | None | FimA<br>PapA<br>SfaA<br>MrkA |
| NTUH-K2044 | *K. pneumoniae* | 23 | KL1 | K1 | O1/O2v2 | O1 | Penicillins (SHV-11) | 4 – aerobactin and/or salmochelin with yersiniabactin (without colibactin) | rmp 1; KpVP-1 (truncated), rmp 3; ICEKp1 (truncated) / rmpA2_3-47% | IncHI1B(pNDM-MAR)/repB_KLEB | FimA<br>PapA<br>SfaA<br>MrkA |
| 664 | *K. pneumoniae* | 37 | KL118 | Unknown | Unknown (OL101) | Unknown (OL101) | Aminoglycosides (aadA2)<br>Cephalosporins (SHV-2A)<br>Phenicol (cmIA4 (homolog))<br>Sulfonamides (sul1, sul2)<br>Trimethoprim (dfrA15 (homolog)) | 0 | -/- | IncFIB(K)<br>Col440I | FimA<br>PapA<br>SfaA<br>MrkA |
| 714 | *K. pneumoniae* | 280 | KL23 | K23 | O1/O2v2 | O2afg | Penicillins (SHV-27) | 0 | -/- | IncFIB(K)<br>IncFII(K) | FimA<br>PapA<br>SfaA<br>MrkA |
| 3557 | *K. pneumoniae* | 20 | KL28 | K28 | O1/O2v2 | O1 | Penicillins (SHV-187) | 0 | -/- | IncFIB(K)<br>IncFII(K) | FimA<br>PapA<br>SfaA<br>MrkA |
| 4773 | *K. pneumoniae* | *80c3 | KL28 | K28 | OL104 | unknown<br>(OL104) | Penicillins (SHV-60) | 0 | -/- | None | FimA<br>PapA<br>SfaA<br>MrkA |
| 4831 | *K. pneumoniae* | 1083 | KL132 | Unknown | O1/O2v1 | O1 | Penicillins (SHV-1)<br>Tetracycline (tet(D)) | 0 | -/- | IncFIB(K) | FimA<br>PapA<br>SfaA<br>MrkA |
| 5022 | *K. pneumoniae* | 628 | KL123 | Unknown | O3b | O3b | Penicillins (SHV-1) | 0 | -/- | IncFIB(K) | FimA<br>PapA<br>SfaA<br>MrkA |
| 5256 | *K. pneumoniae* | 36 | KL102 | Unknown | O1/O2v2 | O2afg | Penicillins (SHV-11) | 0 | -/- | IncFIB(K)(pCAV1<br>099-114)<br>Col(pHAD28) | FimA<br>PapA<br>SfaA<br>MrkA |
| 5712 | *K. pneumoniae* | 45 | KL24 | K24 | O1/O2v1 | O2a | Penicillins (SHV-1) | 1 -<br>yersiniabacti<br>n only | -/- | IncFIB(pKPHS1) | FimA<br>PapA<br>SfaA<br>MrkA |
| 6486 | *K. variicola* | 925 | KL113 | Unknown | O3/O3a | O3/O3a | Penicillins (LEN-2) | 0 | -/- | None | FimA<br>PapA<br>SfaA<br>MrkA |
| 6918 | *K. pneumoniae* | 13 | KL30 | K30 | O1/O2v1 | O1 | Penicillins (SHV-1) | 0 | -/- | IncFIB(K)<br>Col(pHAD28)/C<br>ol440II | FimA<br>PapA<br>SfaA<br>MrkA |
| 7191 | *K. pneumoniae* | 2425 | KL7 | K7 | O3b | O3b | Penicillins (SHV-11) | 0 | -/- | None | FimA<br>PapA<br>SfaA<br>MrkA |
| 7637 | *K. pneumoniae* | 307 | KL102 | Unknown | O1/O2v2 | O2afg | Aminoglycosides (aac(3)-lia, strA, strB)<br>Cephalosporins [3rd Gen(CTX-M-15)]<br>Floroquinolones (qnrB1, GyrA-83I, ParC-80I)<br>Penicillins (TEM-1D, SHV-28)<br>Sulfonamides (sul2)<br>Tetracycline (tet(A))<br>Trimethoprim (dfrA14 (homolog)) | 0 | -/- | Col(pHAD28) | FimA<br>PapA<br>SfaA<br>MrkA |
| 7951 | *K. pneumoniae* | 1380 | KL23 | K23 | Unknown (OL102) | Unknown (OL102) | Penicillins (SHV-26) | 0 | -/- | IncFIB(K)<br>IncFII(K)<br>Col440II | FimA<br>SfaA<br>MrkA |
| 8039 | *K. pneumoniae* | 253 | KL112 | Unknown | O1/O2v2 | O1 | Penicillins (SHV-36) | 0 | -/- | Col(pHAD28)<br>IncFIB(K)<br>IncFII(K)<br>Col440I<br>Col440II<br>IncFIB(pNDM-Mar) | FimA<br>PapA<br>SfaA<br>MrkA |
| 8341 | *K. pneumoniae* | 4073 | KL5 | K5 | O1/O2v1 | O1 | Penicillins (SHV-37) | 0 | -/- | None | FimA<br>PapA<br>SfaA<br>MrkA |
| 8715 | *K. pneumoniae* | 188 | KL122 | Unknown | O1/O2v2 | O2afg | Penicillins (SHV-37) | 0 | -/- | IncFII(K)<br>IncFIB(K)<br>Col440I | FimA<br>PapA<br>SfaA<br>MrkA |
| 10072 | *K. pneumoniae* | 784 | KL146 | Unknown | O1/O2v2 | O1 | Penicillins (SHV-1)<br>Floroquinolones (GyrA-83Y) | 0 | -/- | IncFIB(K)(pCAV1<br>099-114)<br>IncN | FimA<br>PapA<br>SfaA<br>MrkA |
| 10079 | *K. variicola* | 595 | KL16 | K16 | O5 | O5 | Penicillins (LEN-16) | 1 -<br>yersiniabacti<br>n only | -/- | IncFIB(K)(pCAV1<br>099-114) | FimA<br>PapA<br>SfaA<br>MrkA |
| 10081 | *K. variicola* | 616 | KL54 | K54 | O5 | O5 | Penicillins (LEN-9) | 0 | -/- | IncFII(K)<br>IncFIA(pBK3068<br>3)<br>Col440I | FimA<br>PapA<br>SfaA<br>MrkA |
| 10403 | *K.*<br>*quasipneumoniae* | *4562 | KL101 | Unknown | O3/O3a | O3/O3a | Penicillins (OKP-A-11) | 0 | -/- | None | FimA<br>PapA<br>SfaA<br>MrkA |
| 10856 | *K. pneumoniae* | 152 | KL46 | K46 | O3b | O3b | Penicillins (SHV-1) | 0 | -/- | IncFIB(K)<br>INCFIA(pBK306<br>83) | FimA<br>PapA<br>SfaA<br>MrkA |
| 11465 | *K. pneumoniae* | 3096 | KL58 | K58 | O3b | O3b | Aminoglycosides (aac(6')-Ib-cr, strA, strB)<br>Cephalosporins [3rd Gen(CTX-M-15)]<br>Floroquinolones (qnrB1)<br>Penicillins (OXA-1, TEM-1D, SHV-26)<br>Sulfonamides (sul2)<br>Tetracycline (tet(A), tet(B)(homolog), tetR)<br>Trimethoprim (dfrA14 (homolog)) | 0 | -/- | IncI1-I(Alpha)<br>IncFIA(HI1)<br>IncFIB(K)<br>Col(pHAD28) | FimA<br>PapA<br>SfaA<br>MrkA |
| 11469 | *K. pneumoniae* | 253 | KL30 | K30 | O1/O2v2 | O2afg | Penicillins (SHV-36) | 0 | -/- | IncFIB(K)<br>IncFII(K)<br>Col(pHAD28)<br>Col(pHAD28) | FimA<br>PapA<br>SfaA<br>MrkA |
| 11997 | *K. pneumoniae* | 391 | KL30 | K30 | O1/O2v2 | O1 | Aminoglycosides (aac(6')-Ib-cr, strA, strB)<br>Cephalosporins [3rd Gen(CTX-M-15)]<br>Floroquinolones (qnrB1)<br>Penicillins (OXA-1, TEM-1D, SHV-26)<br>Sulfonamides (sul2)<br>Tetracycline (tet(A))<br>Trimethoprim (dfrA14 (homolog)) | 1 - yersiniabactin only | -/- | IncFIB(K)/IncFII(K)<br>ColRNAI | FimA<br>PapA<br>SfaA<br>MrkA |
| 12022 | *K. pneumoniae* | 5142 | KL27 | K27 | O3b | O3b | Penicillins (SHV-187 (homolog)) | 0 | -/- | None | FimA<br>PapA<br>SfaA<br>MrkA |

### Culturing *K. pneumoniae* in urine broadly increases capsule abundance

Recent studies observed that nutrient-limited conditions promotes cKp and hvKp capsule retention compared to rich culture conditions and that hvKp strains increased capsule abundance when cultured in human urine.^21, 32^ Therefore, we sought to determine if cKp strains also increased capsule abundance in nutrient-limited conditions. We cultured the 25 cKp UTI isolates in sterile-filtered pooled human urine or LB medium (**Table 1**).^23–25^ Cell-associated capsule abundance was quantified by measuring uronic acid content in crude capsule extracts.^33^

As predicted, cKp UTI isolates cultured in urine generally had higher capsule abundance relative to LB medium (**Fig. 1A-B**). However, five UTI isolates (Kp4831, Kp5256, Kp5712, Kp8715, and Kp10856) did not have increased capsule abundance in urine (**Fig. 1A**). Regardless of culture condition, hvKp isolates, KPPR1 and NTUH-K2044, produced similar amounts of capsule compared to cKp isolates, reinforcing recent reports that hvKp isolates do not necessarily produce more capsule than cKp isolates (**Fig. S1A**).^34, 35^ In addition, despite urine generally increasing capsule production, all hvKp and cKp strains grew significantly better in LB compared to urine (**Fig. S3**). Thus, increased capsule abundance in urine was not driven by increased growth relative to LB. Altogether, these data support that capsule abundance is elevated in nutrient-limited environmental conditions like urine, although some strain-dependent variations occur.

**Figure 1.**
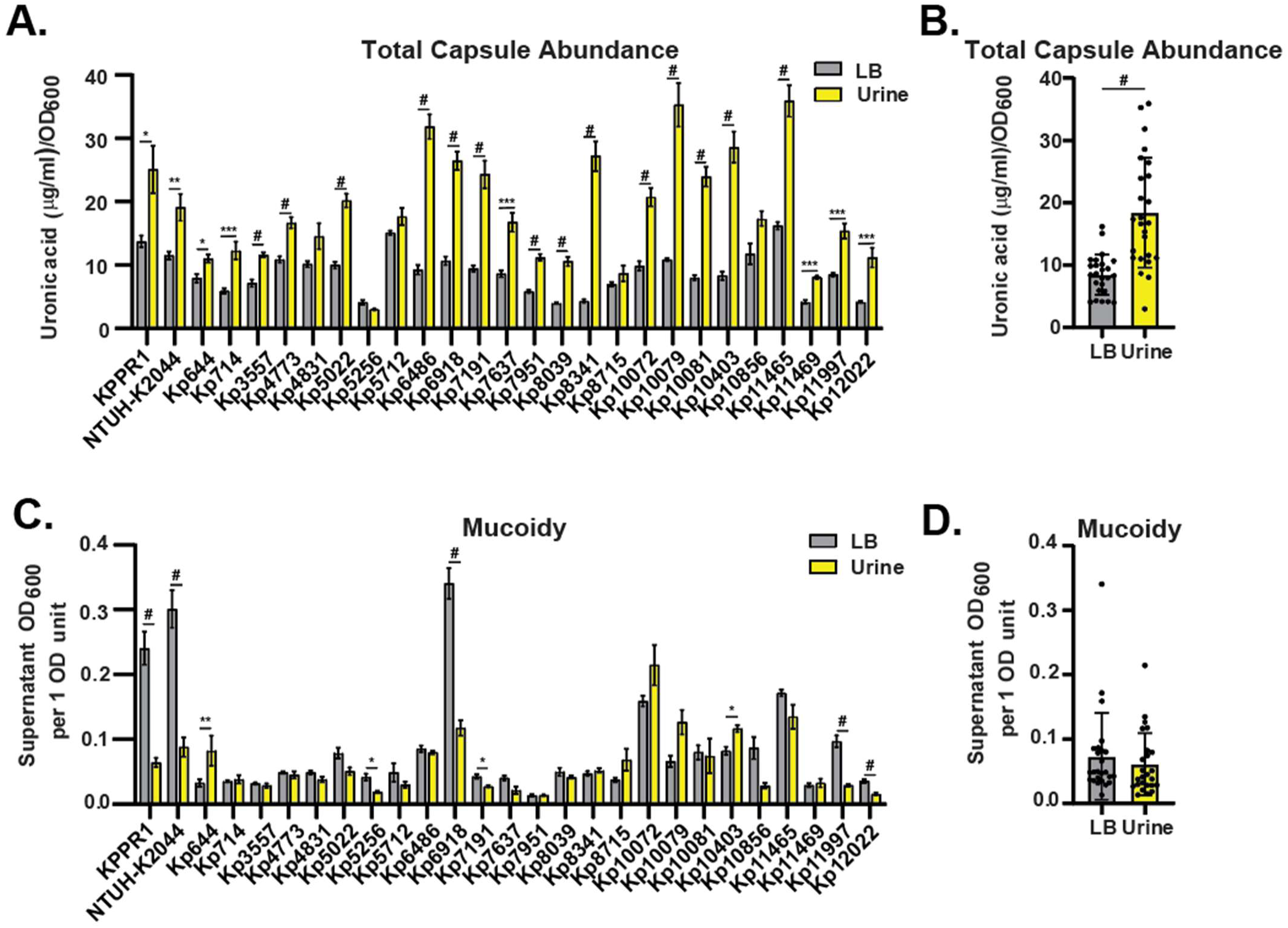
Culturing *K. pneumoniae* in urine increases capsule abundance but exerts heterogeneous effects on mucoidy. Two hvKp strains (KPPR1 and NTUH-K2044) and 25 cKp UTI isolates were cultured in LB medium or in sterile-filtered pooled female human urine. (**A**) Capsule abundance was quantified by measuring uronic acid content in crude capsule extracts. In brief, bacterial cell pellets were washed with PBS, capsule was purified from the cell surface, and then uronic acid content was quantified and normalized to the culture OD_600_. (**B**) Mean capsule abundance of each clinical UTI isolate in **A** was plotted and compared between LB medium and urine. (**C**) Mucoidy was determined by quantifying the supernatant OD_600_ after sedimenting 1 OD_600_ unit of culture at 1,000 *× g* for 5 min. (**D**) The mean sedimentation resistance of each clinical UTI isolate in **C** was plotted and compared between LB medium and urine. (**A** and **C**) Data presented are the mean of experiments performed ≥ 3 independent times, each with biological triplicates. (**B** and **D**) Each dot represents the mean of one strain in **A** or **C**. Error bars represent the standard error of the mean. (**A** and **C**) Statistical significance was determined using an unpaired t test with Holm-Šídák post-test or (**B** and **D**) a paired student’s t-test (* P < 0.05; ** P < 0.01; *** P < 0.001; # P < 0.0001).

### Mucoidy varies between strains and culture conditions

Recent data have established that CPS chain length drives mucoidy, not total capsule abundance.^20, 21, 30, 36^ Furthermore, we have reported that hvKp mucoidy is regulated by sugar and amino acid availability.^37^ ^38^ For example, human urine suppresses mucoidy in hvKp strains.^21^ Interestingly, there have been sporadic reports of cKp strains appearing mucoid in clinical isolates, including from bloodstream infections.^34, 39, 40^ Thus, we hypothesized that some cKp UTI isolates may also be mucoid, and that mucoidy in cKp may also be regulated in response to environmental cues.

We used a standard sedimentation assay to quantify mucoidy, where increased sedimentation resistance indicated increased mucoidy.^33^ Historically, hvKp strains have greater mucoidy than cKp strains, which lack the *rmp* operon.^41^ We found that the hvKp strains, KPPR1 and NTUH-K2044, behaved as expected, exhibiting decreased mucoidy when cultured in urine compared to LB medium (**Fig. S1B**).^21^ Surprisingly, some cKp UTI isolates exhibited mucoidy; however, the cKp UTI isolates did not consistently increase or decrease mucoidy when cultured in human urine relative to LB broth (**Fig. 1C-D**). Notably, none of the cKp strains encode the hypervirulence-associated *rmp* locus or Wzc mutation, both of which are known to induce mucoidy (**Table 1, Fig. S2**).^21, 22, 30^ Thus, some cKp strains likely encode other gene(s) that confer mucoidy and these genes are likely regulated differently than the *rmp* locus in hvKp strains. Finally, comparing mean cKp strain mucoidy to hvKp strains confirmed that cKp strains are generally less mucoid than hvKp strains when cultured in LB medium (**Fig. S1B**).^34^

### Despite encoding adhesins, *K. pneumoniae* UTI isolates rarely hemagglutinate red blood cells

We examined the distribution and homology of common UTI-associated adhesins in our panel of 25 cKp UTI isolates. All isolates encoded homologs to common UTI-associated adhesins: FimA, SfaA, and MrkA (**Table 1**). Furthermore, all isolates encoded a PapA homolog, except Kp7951. Adhesin binding can be detected using hemagglutination assays. Over 70% of UPEC isolates hemagglutinate guinea pig red blood cells (RBCs) via mannose-dependent adhesins like Fim.^42^ Thus, we examined the ability of each strain to hemagglutinate guinea pig RBCs and assessed if adhesin functionality was regulated in response to human urine.

We statically cultured the 25 cKp UTI isolates, two hvKp lab strains, and one UPEC strain, CFT073, in LB medium or urine for 72 hours. Bacteria were suspended in PBS then incubated with guinea pig RBCs for one hour and then assay images were blindly scored.^43^ Three cKp isolates weakly hemagglutinated guinea pig RBCs in a culture-dependent manner (**Fig. 2A**). Although not statistically significant, Kp4831 and Kp11465 only hemagglutinated guinea pig RBCs when cultured in LB medium, while Kp5256 hemagglutinated red blood cells more when pre-cultured in urine (**Fig. 2A**). When comparing all cKp UTI isolates, there was no significant difference in hemagglutination after growth in LB medium versus urine (**Fig. 2B**). Furthermore, no differences in hemagglutination were observed between hvKp or cKp strains in either medium (**Fig. S1C**). Notably, even the cKp UTI isolates that did hemagglutinate guinea pig RBCs, were less effective than UPEC strain, CFT073 (**Fig. 2A**). Altogether these data reveal that *K. pneumoniae* adhesins do not hemagglutinate guinea pig RBCs well, suggesting that cKp adhesins may not be expressed under the tested conditions or may not strongly bind moieties on the guinea pig RBC surface.

**Figure 2.**
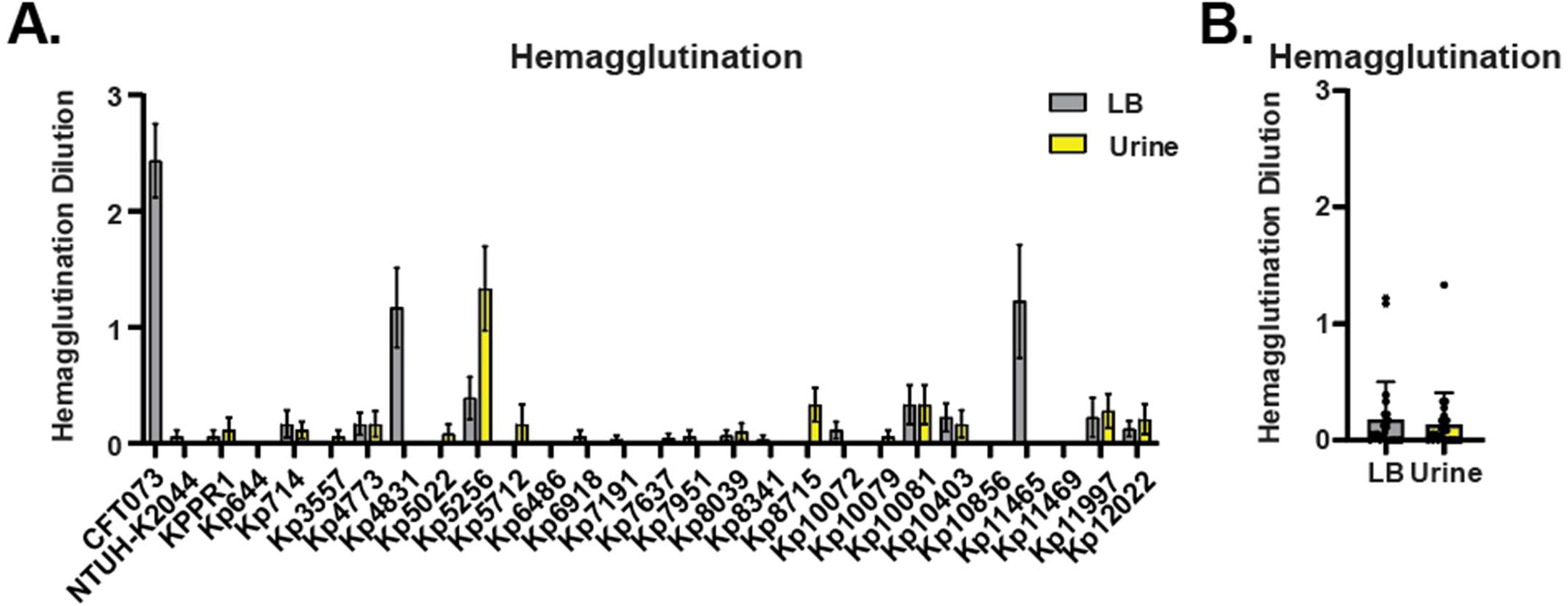
*K. pneumoniae* UTI isolates weakly and rarely hemagglutinate guinea pig red blood cells (RBCs). Twenty-five *K. pneumoniae* clinical UTI isolates and two hvKp laboratory strains, KPPR1 and NTUH-K2044, were cultured in either LB medium or sterile-filtered pooled human urine then serially diluted 1:1 with guinea pig RBCs and incubated to assess hemagglutination. (**A**) The dilution at which hemagglutination occurred was blindly scored and averaged. UPEC strain CFT073 was used as a positive control. Data presented are the mean of experiments performed ≥ 3 independent times, in triplicate. (**B**) Mean hemagglutination of each clinical strain was plotted and compared between LB medium and urine. Statistical significance was assessed using (**A**) an unpaired t test with Holm-Šídák post-test or (**B**) a paired student’s t-test.

### Culturing UTI isolates in urine increases complement resistance

Human serum resistance measures the ability of bacteria to evade complement-mediated bactericidal activity, an important virulence factor for bacteria during UTI.^44, 45^ Both hvKp strains and a sub-set of the cKp UTI isolates were cultured in LB broth or urine prior to human serum exposure to compare the effect of growth environment on complement susceptibility. The subset of clinical isolates (N = 10) represented a variety of mucoidy, capsule abundance, and hemagglutination phenotypes quantified in **Figs. 1–2**. Serum resistance was determined by incubating 2×10^6^ CFU of bacteria in 90% active serum for 90 minutes (**Fig. 3A-B**). Output CFUs were enumerated and normalized to input CFUs. Heat-inactivated serum was tested in parallel as a negative control (**Fig. 3C-D**). A percent survival less than 100% indicates susceptibility to complement-mediated killing.

**Figure 3.**
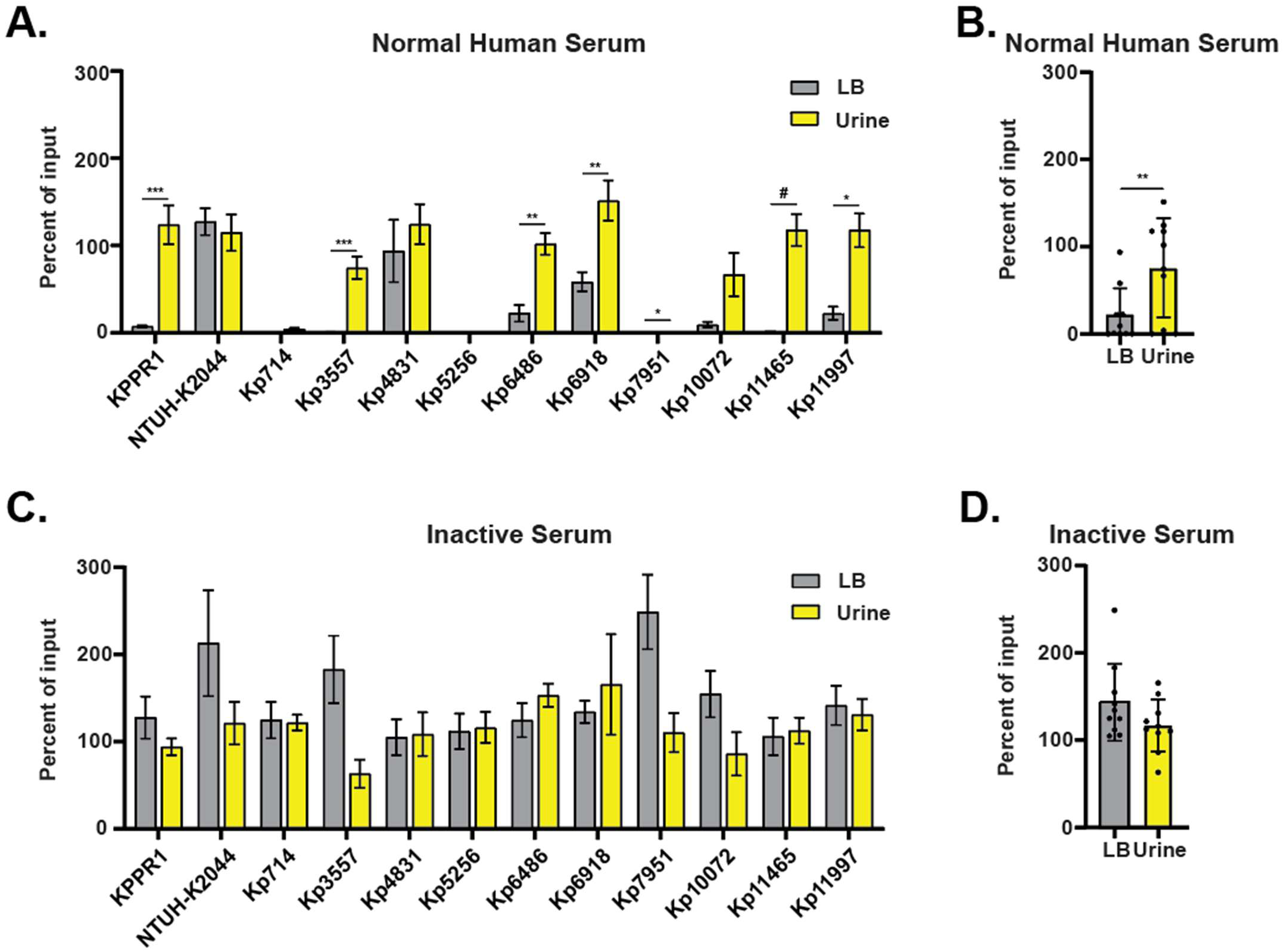
*K. pneumoniae* are more resistant to complement-mediated killing when cultured in urine. Ten *K. pneumoniae* cKp UTI isolates as well as two hvKp laboratory strains, KPPR1 and NTUH-K2044, were cultured in either LB medium or sterile-filtered pooled human urine, washed in PBS, and then 1×10^5^ CFU/mL were co-incubated with (**A-B**) 90% active normal human serum or (**C-D**) 90% heat-inactivated normal human serum at 37 °C for 90 min. The percent survival was calculated by dividing the output CFUs by input CFUs for each strain. Mean serum resistance of each clinical UTI isolate in (**B**) 90% active normal human serum or (**D**) 90% heat-inactivated normal human serum was plotted and compared between LB medium and urine. (**A** and **C**) Data presented are the mean of experiments performed ≥ 3 independent times, each in triplicate. (**B** and **D**) Each dot represents the mean of one strain in **A** or **C**. Statistical significance was determined using (**A** and **C**) an unpaired t test with Holm-Šídák post-test or (**B** and **D**) a paired student’s t-test (* P < 0.05; ** P < 0.01; *** P < 0.001; # P < 0.0001).

Most cKp UTI isolates, as well as KPPR1, exhibited increased complement resistance when cultured in urine compared to LB medium (**Fig. 3B**). NTUH-K2044, Kp714, Kp4831, Kp5256, and Kp10072 did not have significant difference in serum resistance between the two conditions, despite NTUH-K2044, Kp714, and Kp10072 having increased capsule in urine (**Fig. 3A** and **Fig.1A**). NTUH-K2044 and Kp4831 had high survival in both LB and urine, suggesting that some strains are intrinsically more resistant to complement, independent of capsule abundance (**Fig. 3A**). Notably, Kp5256 (likely acapsular), Kp7951, and Kp714 had practically no serum survival in either condition (**Fig. 3A**). Overall, the cKp strains displayed a significant increase in bacterial survival when cultured in urine (**Fig. 3B** and **S3D**). As expected, all isolates survived in heat-inactivated serum regardless of culture condition (**Fig. 3C-D**). Combined, these data indicate that relative to LB medium, culturing cKp UTI isolates in urine increases resistance to complement-mediated killing, likely due to increased capsule abundance.

### Bladder cell association and invasion are driven by strain differences, not environmental conditions

We then modeled cKp bladder cell association and invasion using T24 bladder cells to determine whether strain differences or environmental conditions might impact bladder colonization. From the isolates used in **Fig. 3**., we further down sampled to seven clinical UTI isolates and one hvKp strain, still encompassing a variety of quantified phenotypes (**Fig. 1-3**). Despite invasion occurring at a low frequency, there were some association and invasion differences observed between strains (**Fig. 4C**). Only Kp7951 exhibited a significant difference in T24 bladder cell association or invasion after culturing in urine relative to LB medium (**Fig. 4A** and **4C**). Compared to KPPR1, Kp7951 and Kp6486 both bound bladder cells more after growth in either culture conditions, while Kp4831 only bound bladder cells more after culture in LB medium (**Fig.4A**). Kp7951 also exhibited a significant increase in bladder cell invasion compared to KPPR1 after culture in LB medium (**Fig. 4C**). Although there was a trend toward cKp exhibiting increased in bladder cell association and invasion compared to hvKp, there was no significant difference (**Fig. S1E-F**). Overall, there was no medium-dependent difference in bladder cell association or invasion (**Fig. 4B** and **D**).

**Figure 4.**
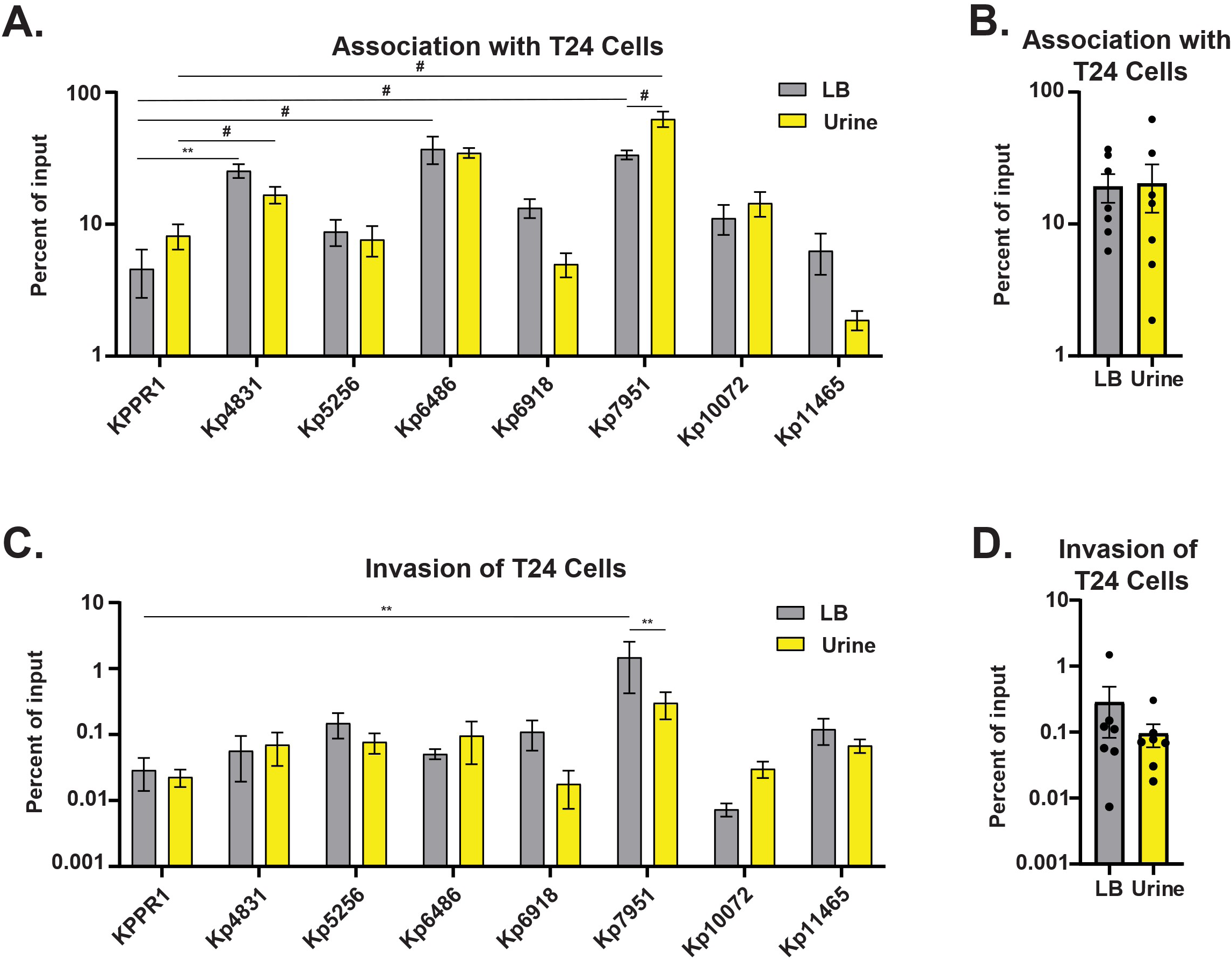
Bladder cell association and invasion are strain-dependent and not influenced by urine. Seven cKp UTI isolates and hvKp strain, KPPR1, were cultured overnight in either LB medium or sterile-filtered pooled human urine. The bacterial cells were washed with PBS and then resuspended in McCoy medium. T24 bladder cells were infected with each strain at an MOI 10 for 2 h, followed by an incubation at 4°C for 1 h (**A-B**) without treatment or (**C-D**) with 100 μg/mL gentamicin. Samples were washed three times with PBS and then lysed with Triton-X100. Input and cell-associated bacterial colony forming units (CFUs) were enumerated on LB agar. (**A** and **C**) Data presented are the log of the mean of experiments performed ≥ 3 independent times, each in triplicate. (**B** and **D**) Each dot represents the mean of one strain in **A** or **C**. Error bars represent the standard error of the mean. Statistical significance was determined using (**A** and **C**) two-way ANOVA with Bonferroni post-test or (**B** and **D**) a paired student’s t-test (* P < 0.05; ** P < 0.01).

### All cKp UTI isolates establish pyelonephritis, despite phenotypic heterogeneity

Four strains representing a variety of phenotypes were selected to assess *in vivo* fitness in a murine UTI model. For example, all four strains significantly increased capsule abundance in urine compared to LB medium, but Kp4831 and Kp11465 hemagglutinated guinea pig RBCs while Kp6918 and Kp10072 did not (**Fig. 1B**). *K. pneumoniae* strains were cultured in either LB or urine, then washed in PBS, and 10^8^ CFUs were transurethrally inoculated into female mice. After 48 hours, bacterial burdens in the bladder, kidneys, spleen, and urine were enumerated. There was no significant difference in bacterial burdens between strains at each site, nor were there significant differences between culture conditions prior to infection (**Fig. 5A-D**). When the data from the four strains were combined, there were no significant differences in median bacterial burdens between host sites (organ or urine) or culture conditions (**Fig. 5E**). However, when the frequency of recoverable CFUs from each host site was quantified, there were significant differences in the percentage of infected organs across the host sites (**Fig. 5F**). Specifically, all *K. pneumoniae* strains colonized kidneys more frequently than either bladders or spleens. Spleens were rarely colonized with bacteria (<15%), suggesting that *K. pneumoniae* infected the kidneys via the ascending route rather than hematogenous spread. These data reveal that despite being phenotypically diverse, the four cKp clinical UTI isolates all successfully colonized the urinary tract, predominantly establishing pyelonephritis.

**Figure 5.**
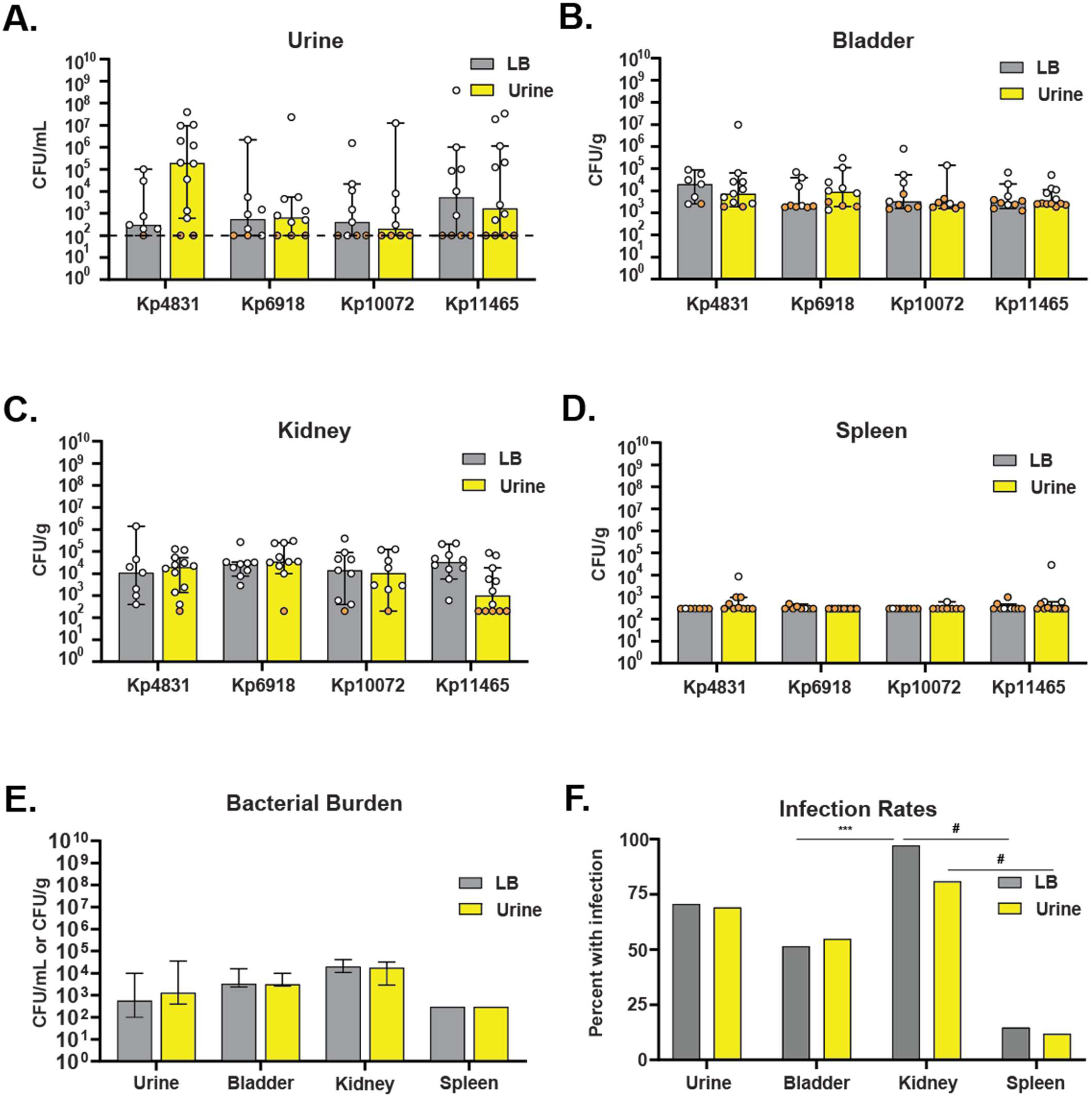
Despite phenotypic heterogeneity, cKp UTI isolates exhibit kidney tropism. Four UTI isolates were cultured in either LB medium or sterile-filtered pooled human urine, then washed and resuspended in PBS prior to transurethral inoculation into female CBA/J mice. After 48 h, the bacterial burdens in the (**A**) urine, (**B**) bladder, (**C**) kidneys, and (**D**) spleens were enumerated. Each dot represents an individual mouse, the dashed line represents the limit of detection (100 CFU/mL), and the orange dots indicate no bacteria were detected in the outputs. (**E**) The combined median bacterial burden of the four strains from **A-D** was calculated for each organ or urine. (**F**) Organs and urine with no recoverable CFUs in **A-D** were classified as ‘uninfected’, and the percent infected was calculated. Infections with each strain were performed on at least two independent days. Statistical significance was determined using two-way ANOVA with Tukey’s post-test (*** P < 0.001; # P < 0.0001).

### Capsule abundance and serum resistance are correlated

Using a correlation analysis matrix, we tested if there were relationships between the various virulence phenotypes by combining data collected for all strains in both culture conditions (**Fig. 6**). We only observed a significant correlation between capsule abundance and serum resistance (P = 0.0002). These data align with prior observations that increased capsule presence promotes complement resistance.^46, 47^ While not significant, mucoidy tended to correlate with T24 association (P = 0.09858), which aligns with prior reports that mucoidy interferes with bacterial binding to host cells.^34, 36^ Interestingly, hemagglutination overall had very low correlation with other virulence factors. Thus, most quantified phenotypes did not appear to be related to each other, except that elevated capsule production resulted in greater protection from complement-mediated killing.

**Figure 6.**
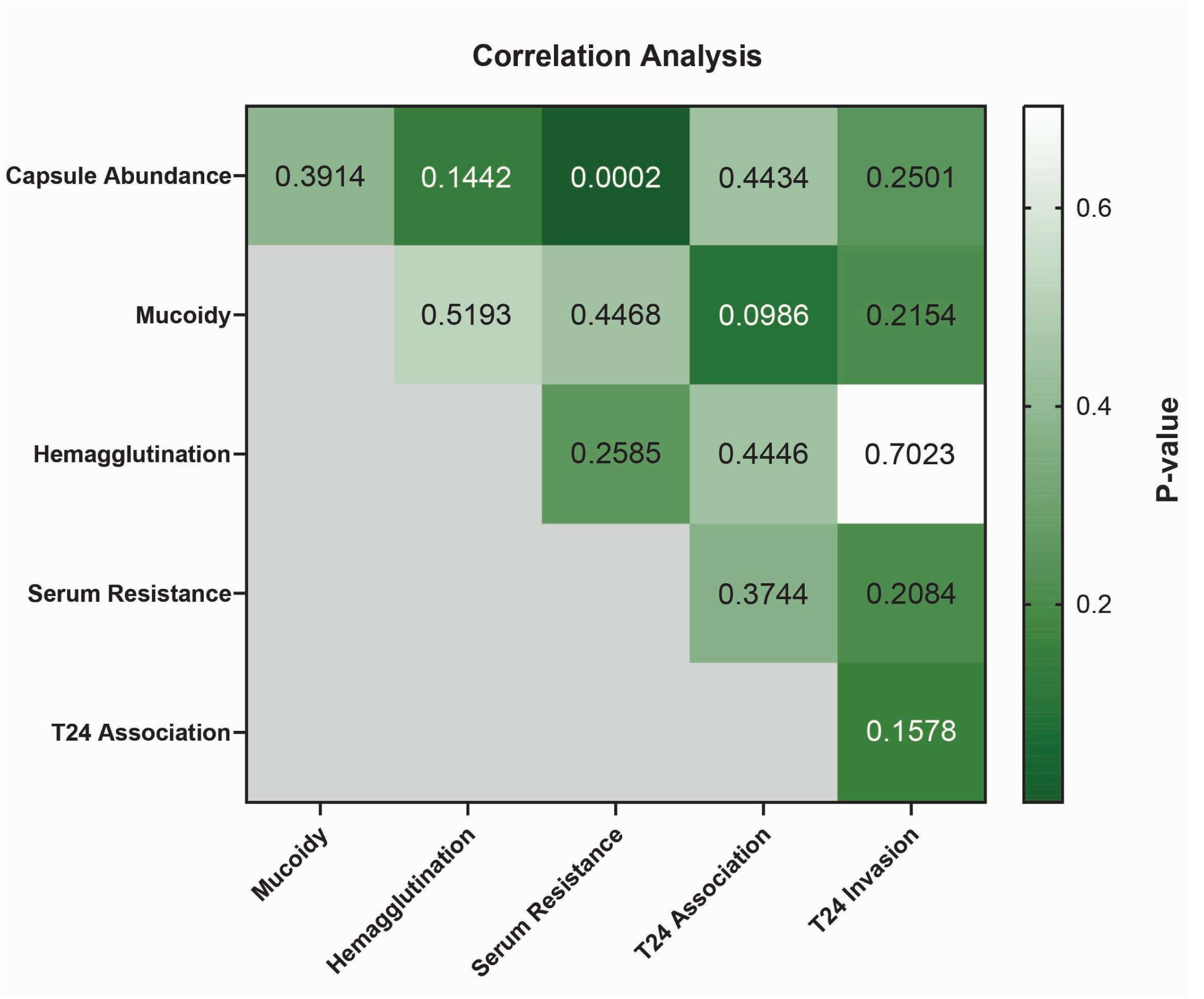
Correlation analyses between quantified *K. pneumoniae* fitness phenotypes. A correlation matrix was used to compare relations between the virulence phenotype quantified in Fig. 1-4 using a nonparametric Spearman correlation and two-tailed p values with a 95% CI. The p values are displayed, and p < 0.05 was deemed significant. The lighter shade indicates less significant correlation, while the darker green indicates more significant correlation.

## DISCUSSION

Despite being the second leading cause of UTIs, *K. pneumoniae* uropathogenesis is understudied relative to UPEC. Although *K. pneumoniae* employs some of the same virulence factors as UPEC, it relies more heavily on capsule for virulence and more frequently exhibits mucoidy. We found that while some fitness-associated phenotypes were differentially expressed based on environmental cues, many phenotypes were expressed in a strain-dependent manner, *i.e.,* there were no consistent differences between growth media, rather phenotypes differed between strains. Despite the observed heterogeneity amongst *K. pneumoniae* clinical UTI isolates, strains were able to establish UTIs *in vivo,* regardless of virulence phenotype expression. Interestingly, *K. pneumoniae* strains all had robust kidney colonization, suggesting an unknown conserved bacterial property that promotes UTI ascension.

To examine the genomic properties encoded by *K. pneumoniae* clinical UTI isolates, we analyzed 25 clinical UTI isolates using Pathogenwatch and performed BLAST analyses with known adhesin sequences.^31^ The *K. pneumoniae* clinical UTI isolates had high adhesin homology to UPEC, with all but one strain encoding FimA homologs with 81-83% identity, PapA homologs with 25-31% identity, SfaA with 57-65% identity homologs, and MrkA homologs with 93-95% identity (**Table 1**). The other strain, Kp7951 encoded all homologs to a lesser degree, but did not encode a PapA homolog. Thus, *K. pneumoniae* encodes many of the same adhesins as UPEC, including FimA, which primarily mediates bladder cell adherence and invasion via type 1 pili.^48, 49^ In agreement with our adhesin bioinformatic analysis, we found that *K. pneumoniae* clinical isolates successfully adhered to and invaded T24 bladder cells, with strain-dependent differences (**Fig. 4**).^48^

CPS are produced by both UPEC and *K. pneumoniae* and block innate immune responses, such as phagocytosis and complement-mediated killing (**Fig. 1A**).^50, 51^ Serum resistance, which allows UPEC to evade complement-mediated killing, is a key virulence determinant for urosepsis.^45^ *K. pneumoniae*, like UPEC, also demonstrates serum resistance in a strain-dependent manner, with only three strains exhibiting little to no complement resistance (**Fig. 3A**). While both *E. coli* and *K. pneumoniae* produce CPS, *K. pneumoniae* controls CPS length and uniformity to generate a mucoid phenotype, which is an important virulence factor associated with invasive hvKp infections.^20, 21, 30, 36^ Hypermucoid phenotypes have previously been associated with hvKp isolates and more severe infections, but cKp strains can exhibit this phenotype as well.^21, 34, 40^ *K. pneumoniae* clinical UTI isolates produced varying levels of mucoidy, which did not correlate with culture conditions (**Fig. 1C**).

Hemagglutination is mediated by both type 1 and P fimbriae, with type 1 and P fimbriae being attributed to primarily supporting cystitis and pyelonephritis, respectively.^52–56^ UPEC expressing adhesins are often detected using hemagglutination assays.^42^ Type 1 fimbriae undergo mannose-sensitive hemagglutination, while P fimbriae can be detected by mannose-resistant hemagglutination.^57, 58^ Hemagglutination of guinea pig erythrocytes is mediated by type 1 fimbriae and are mannose-sensitive, while hemagglutination of human erythrocytes requires both mannose-sensitive and -resistant mechanisms.^42, 57–59^ More virulent strains of UPEC are more likely to be inhibited by mannose; therefore, we tested *K. pneumoniae* clinical UTI isolates using a mannose-sensitive hemagglutination assay with guinea pig RBCs to determine whether, like UPEC, hemagglutination can be used to predict virulence.^58^ Despite all isolates encoding a FimA homolog, most *K. pneumoniae* clinical UTI isolates demonstrated poor guinea pig RBC hemagglutination (**Fig. 2A**). Similar to UPEC, *K. pneumoniae* encodes FimH, an important adhesin at the tip of type 1 pili.^16^ However, *K. pneumoniae* FimH has lower affinity for mannose than UPEC FimH and therefore hemagglutinates guinea pig RBCs poorly.^16, 60^ These interspecies differences in FimH-mannose affinity align with our observation that the UPEC strain CFT073 hemagglutinated guinea pig RBCs significantly better than all the *K. pneumoniae* strains (**Fig. 2A**).

In summary, while *K. pneumoniae* and UPEC isolates share several virulence mechanisms, including fimbrial adhesins, bladder cell invasion and adhesion, CPS production and serum resistance, they differ in other key aspects. Such that *K. pneumoniae* isolates exhibits mucoidy more frequently than UPEC (though still in a minority of UTI isolates) and lacks robust guinea pig RBC hemagglutination, whereas UPEC demonstrates strong hemagglutination but is rarely mucoid. These distinct virulence profiles suggest that *K. pneumoniae* and UPEC employ different pathogenic strategies during UTI.

In addition to observing phenotypic heterogeneity between the clinical UTI isolates, we also considered how culture media alters *K. pneumoniae* virulence phenotypes, and whether those changes affect uropathogenesis. Despite previous research demonstrating that hvKp strains exhibit decreased mucoidy in human urine compared to LB medium, mucoid cKp isolates did not show a significant difference between the two culture conditions (**Fig. 1C-D**).^21^ Moreover, the different culture conditions did not affect RBC hemagglutination or T24 cell invasion or association (**Fig. 2 and Fig. 4**). One limitation is that some clinical isolates alter their mucoidy in response to the bladder cells culture medium, McCoy’s medium, which may confound the bladder association data. However, as previously reported, capsule abundance tended to be elevated in strains cultured in urine compared to LB medium (**Fig. 1B**).^21^ The increased capsule production is likely driven by the nutrient-limited conditions in urine, which has been reported elsewhere.^21, 32^ Capsule also protects bacteria from complement-mediated killing.^46, 47^ Thus, as expected, serum resistance increased when strains were cultured in urine compared to LB medium (**Fig. 3B**). We then employed a correlation matrix analysis to test the relationship between virulence phenotypes (**Fig. 6**). As expected, capsule abundance and serum resistance significantly correlated. This aligns with prior reports that increased capsule abundance protects from both phagocytosis and complement-mediated killing during an infection.^46, 47^

Despite displaying heterogeneous phenotypes across clinical UTI isolates, *K. pneumoniae* consistently colonized the urinary tract in our murine UTI model. Across the tested organs and urine samples, there was no significant difference in bacterial burdens between strains, organs, or culture condition (**Fig. 5A-E**). However, in both media conditions, the clinical UTI isolates colonized kidneys more frequently than bladders or spleens (**Fig. 5F**). We interpret the lack of spleen colonization to indicate that the bacteria reach the kidneys by ascending the ureters from the colonized bladder, rather than through hematogenous dissemination. Our *in vivo* murine data are consistent with clinical observations that *K. pneumoniae* is more likely to cause severe acute pyelonephritis and urosepsis compared to UPEC.^61^ Additionally, UTI patients infected with *K. pneumoniae* had a significantly higher 30-day mortality rate compared to patients infected with UPEC.^61^ Our data, combined with clinical observation, indicates that *K. pneumoniae* exhibit kidney tropism.^61^ Since this was observed in genetically and phenotypically distinct strains, this suggests that core genomic properties may drive kidney tropism. Further investigation is merited to identify the bacterial and host determinants driving *K. pneumoniae-*kidney interactions and their impact on UTI severity. Understanding these host-pathogen interactions could reveal new intervention points that could be targeted to improve outcomes for patients experiencing a *K. pneumoniae-*incited UTI.

Characterizing this collection of cKp UTI isolates has revealed that *K. pneumoniae* uropathogenesis is likely distinct from UPEC. The data presented here demonstrate that clinical *K. pneumoniae* UTI isolates exhibit heterogeneous expression of virulence phenotypes, some of which are regulated by environmental cues. Yet, despite this heterogeneity, the strains consistently colonize the urinary tract, particularly the kidneys. These observations lead us to propose that core genome elements support *K. pneumoniae* urinary tract ascension, establishing pyelonephritis. We propose that the core genomic elements that support ascension are constitutively expressed or activated in the urinary tract. Future research defining the bacterial and host elements required for ascension to the kidneys will advance our understanding of *K. pneumoniae* uropathogenesis and may present the opportunity for improved UTI management.

## METHODS

### Strains and Culture Conditions

All strains described in these studies are detailed in **Table 1**. Bacteria were cultured in either pooled healthy female human urine or lysogeny broth (LB, 5 g/L yeast extract, 10 g/L tryptone, 0.5 g/L NaCl) at 200 rpm and 37°C in an aerated culture tube at 60° angle, unless otherwise noted.^62^ Solid medium was prepared by adding 20 g/L bacto-agar prior to autoclaving. Urine was pooled from at least five independent donors and then vacuum filter sterilized through a 0.2 µm PES membrane. Sterilized urine was stored in aliquots at −20°C with working volumes stored at 4°C.

### PathogenWatch

*K. pneumoniae* strains were characterized based on genomic sequence as published.^62^ In brief, raw sequence reads of each strain from the NCBI database were assembled into whole genomes using PATRIC (**Table S1**). The genomes for KPPR1 and NTUH-K2044 were obtained directly from the NCBI database. The assembled genomes were uploaded to Pathogenwatch for species identification and characterization.^29, 63, 64^ Adhesin sequences for FimA (c5393, C5011), PapA (c3592, c5188, C4895), and SfaA (C1109) were obtained from uropathogenic *Escherichia coli* (UPEC) strains *E. coli* O6:K2:H1 CFT073 (ECC) and *E. coli* O18:K1:H7 UTI89 (ECI) using the Kyoto Encyclopedia of Genes and Genomes (KEGG) database.^31^ An adhesin sequence for MrkA (protein id: ABW83989.1) was obtained from UPEC strain MS2027.^65^ Amino acid homology for each adhesin was compared to each *K. pneumoniae* strain using tBLASTn (**Table S2**). Those proteins with ≥25% identity were classified as adhesin-positive in **Table 1**.

### Wzc alignments

The Wzc protein alignment was performed as described in.^21^ In brief, raw sequence reads of Kp10072, Kp10079, Kp10403, Kp10856, Kp11465, Kp664 and Kp6918 from the NCBI SRA database were assembled using PATRIC (**Table S1**).^66^ The *wzc* ORF was identified by searching the assembled genomes for “tyrosine-kinase protein”. The predicted *wzc* ORF was confirmed by a BLAST search to confirm the ORF contained a tyrosine kinase. The ORFs were translated and aligned using Clustal Omega. The alignment was visualized using Jalview.

### Sedimentation assay

Sedimentation assays were used to quantify mucoidy of overnight samples as described with the following modifications.^21^ In brief, overnight samples equivalent to 1 OD_600_ unit of bacteria were transferred to individual 2 mL centrifuge tubes for samples incubated in LB broth and 10 mL centrifuge tubes for samples incubated in urine if the total amount of urine exceeded 2 mL. The samples were centrifuged at maximum speed for 15 min. All but 50 µL of supernatant fluid was decanted. The pellet was resuspended in 1 mL of PBS and centrifuged at 1,000 *x g* for 5 min. The optical density at 600 nm (OD_600_) of the supernatant was measured to determine the sedimentation resistance.

### Uronic acid quantification

Capsule abundance was quantified based on uronic acid content in CPS isolated from overnight cultures as previously described.^21^ In brief, overnight cultures containing 1 OD_600_ unit were transferred to individual 2 mL centrifuge tubes for strains grown in LB and 10 mL centrifuge tubes for strains grown in urine. The samples were washed to remove uronic content present in urine by centrifuging at maximum speed for 15 min. All but 50 µL of supernatant fluid was decanted. The pellet was resuspended, washed in 1 mL of PBS, and centrifuged at maximum speed for 15 min. After centrifugation, 750 µL of supernatant fluid was removed by pipette. Then, 50 µL of 1% Zwittergent 3-14 in 100 mM citric acid was mixed into the sample and the pellet was resuspended by pipetting. Hereafter, capsule was purified as previously described.^21^ In brief, samples were incubated at 50°C for 20 min and then centrifuged at high speed for 5 min. Capsular polysaccharides were precipitated with 80% ethanol on ice, centrifuged, and the pellets rehydrated in deionized water. Uronic acid content was quantified using 3-hydroxydiphenyl as described.^36, 67^

### Growth Curves

Growth curves were generated using the protocol described.^62^ In brief, bacterial strains were cultured overnight in triplicate in 3 mL of LB medium. The cultures were normalized to an OD_600_ of 0.001 in LB medium or urine, and then 100 µL was aliquoted into a microplate. The OD_600_ was recorded on a Biotek plate reader every 30 min for 16 h at 37°C with continuous, orbital shaking [282 rpm (3 mm)].

### Hemagglutination

Hemagglutination protocol was performed as described.^43^ Bacteria were statically cultured in either 3 mL of LB or pooled urine for 72 h at 37 °C. 400 μL (LB) or 2 mL (urine) of culture were centrifuged at 5,000 *x g* for 5 min and cells were gently washed in 300 μL of PBS. The cell suspensions were serially diluted 1:1 with PBS in round-bottom 96-well plates. 200 μL of guinea pig erythrocytes (Innovative Research: IGPRBC10ML) were pelleted at 5,000 *x g* for 5 min and washed 3 times in 1 mL PBS. Erythrocytes were resuspended in 15 mL of PBS (final concentration 1% vol/vol) and 150 µL were distributed to each well. Bacteria and erythrocytes were gently mixed and incubated statically at room temperature for 1 h prior to imaging. To quantify, hemagglutination was blindly scored by three individuals. The dilution at which individuals identified hemagglutination was averaged for each round.

### Serum Resistance

This assay was performed as described with the following adaptations.^36^ 1 mL of each overnight culture was centrifuged at 16,000 *x g* for 10 min and the pellet resuspended in 1 mL of PBS. The samples were normalized to an OD_600_ of 0.002 in 200 μL PBS to achieve a stock of 2×10^6^ CFU/mL. This working stock was then diluted 1:10 into 96-well plates with either pooled normal human complement serum (Innovative Research, CAT: ICSER50ML) or heat-inactivated (1 h at 58 °C) normal human serum and incubated at 37°C for 90 minutes. Input and output CFU/mL were serially diluted 1:10 with PBS and spot plated on LB agar. Plates were incubated at 30°C until CFUs were visible. To quantify serum resistance, CFUs were averaged and the percent survival relative to input was plotted.

### Bladder cell association and invasion

Cell association and invasion assays were performed using immortalized T24 bladder cells derived from the bladder of a female with colorectal adenocarcinoma (ATCC HTB-4). The cells were maintained in McCoy’s 5A medium with L-glutamine, sodium bicarbonate supplemented with 10% heat-inactivated fetal calf serum (Corning), 100 U/mL penicillin, and 100 µg/mL streptomycin in an atmosphere of 5% CO_2_. Confluent cells (∼8 × 10^4^ cells/well) in 24-well tissue culture-treated dishes were washed three times with 1 mL of PBS, and then 1 mL of 8 × 10^5^ CFU/mL bacteria (MOI 10) in additive-free McCoy’s 5A medium was added to each well. Samples were spun at 54 *× g* for 5 min and then incubated at 37°C, 5% CO_2_ for 4 h, followed by an incubation at 4°C for 1 h with or without 100 μg/mL gentamicin. Samples were washed three times with PBS and then lysed with 1 mL of 0.4% Triton-X100 in PBS for 8 min on a rotary shaker. Input and cell-associated bacterial counts were determined by serial dilution and CFU enumeration on LB agar.

### Murine model of ascending urinary tract infection

These studies were performed and adapted as described. ^43, 68–70^ In brief, female CBA/J mice (6 to 8 weeks old) from Jackson Laboratories were transurethrally inoculated with 50 μL of bacterial suspension (1 × 10^8^ CFU) delivered over 30 s using a sterile pediatric intravenous-access cannula (divested of its needle) connected to an infusion pump (Harvard Apparatus). The procedure was performed under anesthesia with ketamine/xylazine, and mice were recovered on a heated pad. Forty-eight hours post-inoculation, urine was collected and then mice were euthanized by isoflurane overdose. Bladders, kidneys, and spleens were aseptically removed and homogenized according to the manufacturer’s instructions (Bullet Blender Gold, Next Advance). Urine and organ homogenates were serially diluted in PBS and enumerated on LB agar.

### Ethics statement

The animal protocol was approved by the Institutional Animal Care and Use Committee (IACUC) at the University of Toledo (#400104-UT) and research was performed accordingly. Urine collection at the University of Toledo was approved by and performed in accordance with the Institutional Review Boards (IRB) of the University of Toledo (#301539-UT). In brief, female human urine was collected from healthy volunteers at the University of Toledo Health Science Campus who were not diabetic, not menstruating, not pregnant, and had not taken antibiotics within two weeks prior to donating. Written informed consent was received from participants prior to inclusion in the study. These protocols are compliant with the guidelines established by the National Institutes of Health for animal and human subject research.

### Statistics

All *in vitro* experiments include data collected from three biological replicates performed in three independent experiments. All *in vivo* experiments were performed at least twice per strain. All statistical analyses were computed in Prism 10 (GraphPad Software, La Jolla, CA). For experiments comparing multiple groups, significance was calculated using two-way ANOVA with a Bonferroni post-test to compare specific groups. A one sample t-test was applied when comparing two groups or a single group to a hypothetical value of 1.00. A multiple correlation analysis was used to compare quantifiable virulence factors throughout several experiments. In all instances, results were considered significant if the P value was less than or equal to 0.05.

## ACKNOWLEDGEMENTS

The authors thank Dr. Michael Bachman, Dr. Jay Vornhagen, and Sophia Rose Mason for providing clinical isolates. We would like to thank Dr. R. Mark Wooten, Dr. John Presloid, Dr. Chelsie Armbruster, and Brian Learman for technical assistance. We would like to acknowledge Saroj Khadka and Emily Kinney for blinded scoring of hemagglutination data and Krista Pettee for cultivating T24 bladder cells. We thank members of the Mike lab for critical feedback, especially Saroj Khadka, Emily Kinney, Dr. Zach Resko, and Drew Stark. Research reported in this publication was supported by the University of Pittsburgh (LAM) and the University of Toledo Research Innovation Program (LAM) and Medical Student Research Program (DAP and ZM).

---

**Figure S1.**
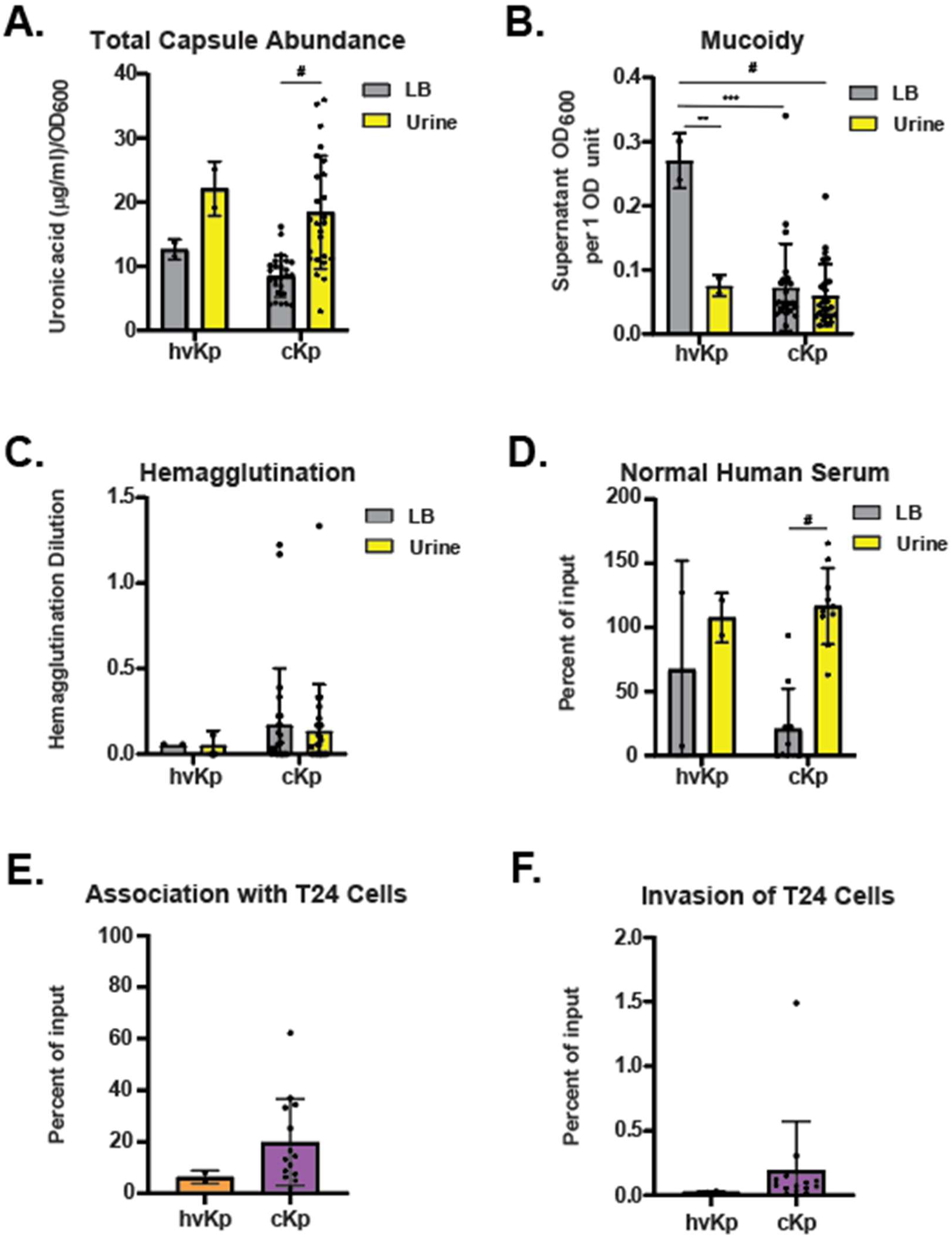
Phenotypic comparisons between hvKp and cKp strains. Values from data presented in the main figures were plotted and stratified by *K. pneumoniae* pathotype. Each dot represents the mean of a single hvKp or cKp strain. Phenotypic differences between growth in sterile-filtered pooled human urine and LB were compared for (**A**) cell-associated capsule abundance, (**B**) mucoidy, (**C**) hemagglutination, and (**D**) human serum resistance. Due to limited data, urine and LB data were combined and compared between hvKp and cKp for (**E**) T24 cell association and (**F**) T24 cell invasion. Each dot represents the mean of one strain and error bars represent standard error of the mean. Statistical significance was determined using (**A-D**) one-way ANOVA with Tukey’s post-test or (**E-F**) Mann-Whitney U test (* P <0.05; ** P < 0.01; *** P < 0.001; # P < 0.0001).

**Figure S2.**
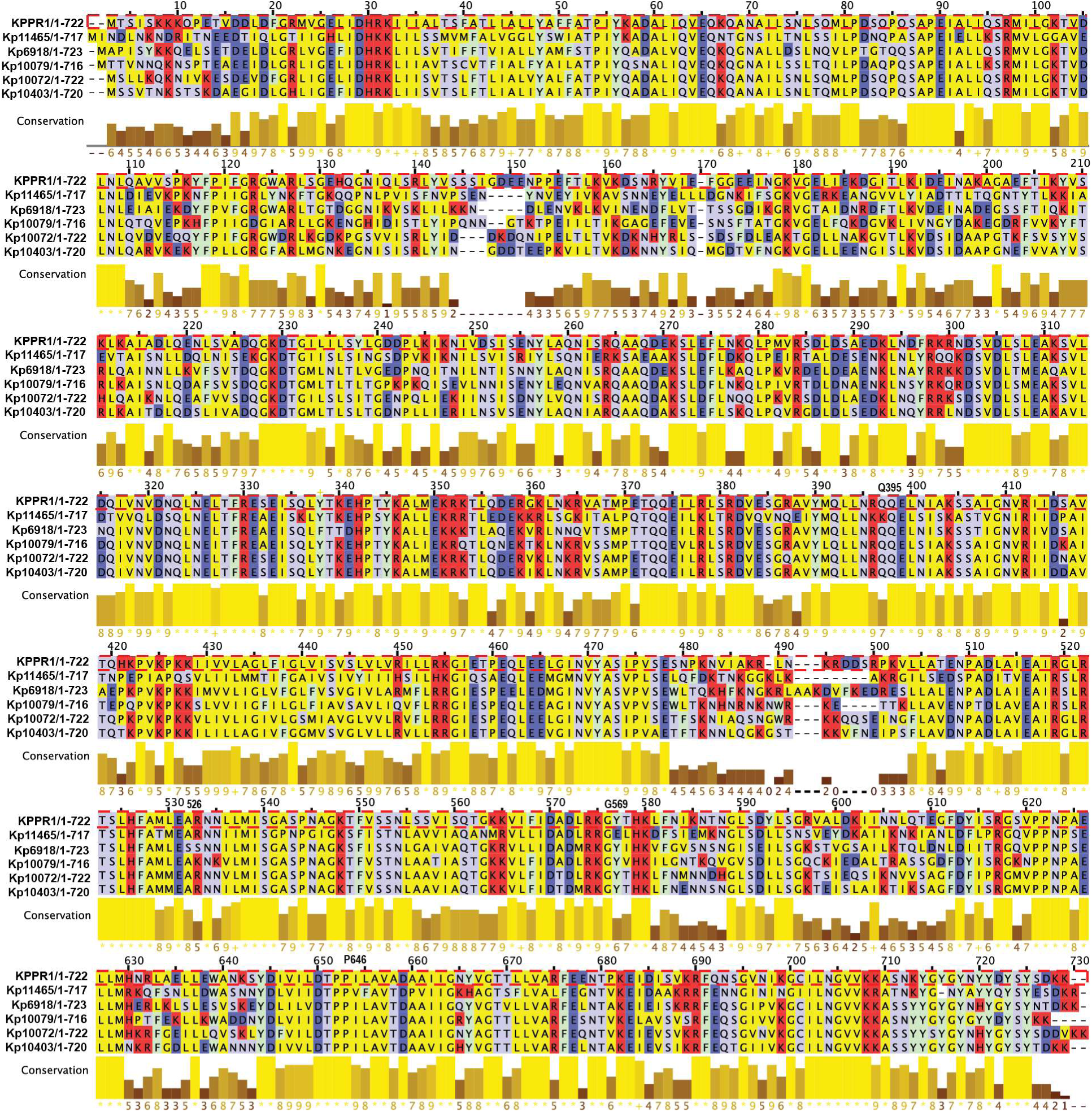
Wzc alignment from mucoid cKp strains. Predicted open reading frames (ORFs) of *wzc* were obtained from unassembled genomes available in the NCBI Sequence Read Archive (SRA). Subsequently, an alignment of the translated ORFs was created using Clustal Omega, and the results were visualized using the Jalview application and Illustrator. In the visualization, the color scheme was employed to represent different types of amino acids: red for basic, dark blue for acidic, light blue for polar uncharged, green for aromatic, and yellow for aliphatic residues. The red dotted line highlights the KPPR1 alignment. The conservation of these amino acid sequences across different strains is depicted through histograms positioned below the alignment.

**Figure S3.**
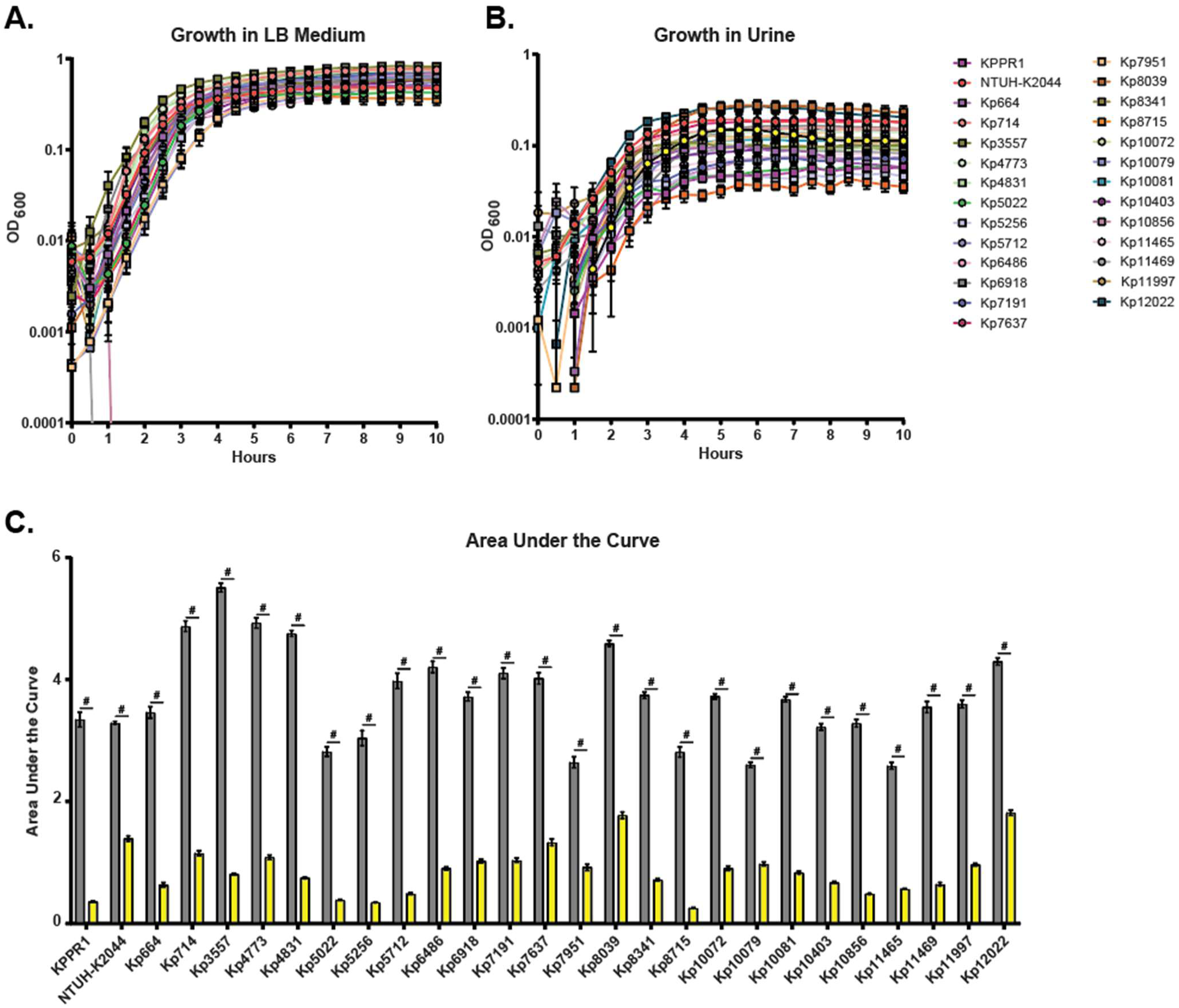
*K. pneumoniae* achieves a lower growth density in urine relative to LB. All strains were cultured in (**A**) LB medium and (**B**) sterile-filtered pooled human urine and then back-diluted to an OD6_00_ 0.001 in the respective medium. Cultures were incubated with shaking for 10 h at 37 ⁰ C and the density measured every 30 minutes. All growth assays were performed ≥ 3 independent times, in triplicate. (**C**) The data presented are the mean of the area under the curve and error bars represent standard error of the mean. Statistical significance was determined using student’s t-test (# P <0.0001).

**Supplemental Table 1.** SR accession numbers for *Klebsiella* UTI isolates. KPPR1 and NTUH-K2044 genomes were preassembled in the NCBI database. The rest of the strains were assembled using BV-BRC genome assembly.

| Strain | Accession Number |
| --- | --- |
| KPPR1 | GCA_000742755.1 |
| NTUH-K2044 | GCA_000009885.1 |
| 664 | SRS1664169 |
| 714 | SRS1664168 |
| 3557 | SRS12803755 |
| 4773 | SRS12904747 |
| 4831 | SRS12904751 |
| 5022 | SRS12904764 |
| 5256 | SRS12904775 |
| 5712 | SRS12904710 |
| 6486 | SRS12803797 |
| 6918 | SRS12803740 |
| 7191 | SRS12803745 |
| 7637 | SRS12803752 |
| 7951 | SRS12803757 |
| 8039 | SRS12803761 |
| 8341 | SRS12803776 |
| 10072 | SRS12902321 |
| 10079 | SRS12902234 |
| 10081 | SRS12902235 |
| 10403 | SRS12902267 |
| 10856 | SRS12902291 |
| 11465 | SRS12902311 |
| 11469 | SRS12902313 |
| 11997 | SRS12902325 |
| 12022 | SRS12902327 |

**Supplemental Table 2.** Percent identity between uropathogenic *E. coli* and *Klebsiella* adhesins. Each number indicates percent identity between the adhesin sequences of the indicated genome.

| *Klebsiella*<br>strain | <u>UPEC CFT073</u> |  |  | <u>UPEC UTI89</u> |  |  | <u>UPEC</u><br>(MS2027) |
| --- | --- | --- | --- | --- | --- | --- | --- |
|  | FimA<br>(c5393) | PapA<br>(c3592) | PapA2<br>(c5188) | FimA<br>(C5011) | PapA<br>(C4895) | SfaA<br>(C1109) | MrkA |
| KPPR1 | 81 | 31 | 32 | 83 | 0 | 57 | 95 |
| NTUH-K2044 | 81 | 34 | 31 | 83 | 0 | 57 | 94 |
| Kp664 | 81 | 41 | 37 | 83 | 39 | 57 | 95 |
| Kp714 | 81 | 40 | 36 | 83 | 39 | 57 | 94 |
| Kp3557 | 81 | 34 | 31 | 83 | 0 | 57 | 95 |
| Kp4773 | 81 | 40 | 36 | 83 | 39 | 60 | 95 |
| Kp4831 | 81 | 40 | 36 | 83 | 39 | 57 | 94 |
| Kp5022 | 81 | 34 | 31 | 83 | 0 | 57 | 94 |
| Kp5256 | 81 | 34 | 31 | 83 | 0 | 57 | 95 |
| Kp5712 | 81 | 34 | 31 | 83 | 0 | 57 | 95 |
| Kp6486 | 81 | 33 | 33 | 82 | 0 | 65 | 93 |
| Kp6918 | 81 | 34 | 31 | 83 | 0 | 57 | 94 |
| Kp7191 | 81 | 34 | 31 | 83 | 0 | 57 | 95 |
| Kp7637 | 81 | 40 | 36 | 83 | 39 | 57 | 95 |
| Kp7951 | 30 | 0 | 0 | 32 | 0 | 34 | 26 |
| Kp8039 | 83 | 34 | 31 | 83 | 0 | 60 | 94 |
| Kp8341 | 81 | 34 | 31 | 83 | 0 | 64 | 95 |
| Kp8715 | 81 | 34 | 31 | 83 | 0 | 64 | 94 |
| Kp10072 | 81 | 34 | 31 | 83 | 0 | 60 | 94 |
| Kp10079 | 81 | 35 | 32 | 83 | 0 | 65 | 93 |
| Kp10081 | 81 | 33 | 25 | 82 | 25 | 65 | 93 |
| Kp10403 | 81 | 34 | 32 | 83 | 26 | 63 | 94 |
| Kp10856 | 81 | 34 | 31 | 83 | 0 | 57 | 94 |
| Kp11465 | 81 | 40 | 36 | 83 | 39 | 57 | 95 |
| Kp11469 | 81 | 34 | 31 | 83 | 0 | 60 | 95 |
| Kp11997 | 81 | 34 | 31 | 83 | 0 | 57 | 94 |
| Kp12022 | 81 | 34 | 31 | 83 | 0 | 57 | 94 |

**Supplemental Table 3.** Growth characteristics of strains in LB and urine.

| Strain | LB |  |  | Urine |  |  |
| --- | --- | --- | --- | --- | --- | --- |
|  | Doubling Time | Area Under the Curve | Endpoint OD | Doubling Time (h) | Area Under the Curve | Endpoint OD<sub>600</sub> |
| KPPR1 | 0.55 | 3.35 | 0.683 | 1.46 | 0.38 | 0.058 |
| NTUH-K2044 | 0.83 | 3.29 | 0.474 | 0.92 | 1.4 | 0.182 |
| Kp664 | 0.63 | 3.47 | 0.683 | 1.05 | 0.64 | 0.058 |
| Kp714 | 0.81 | 4.87 | 0.474 | 1.08 | 1.16 | 0.151 |
| Kp3557 | 0.96 | 5.51 | 0.505 | 1.11 | 0.81 | 0.102 |
| Kp4773 | 0.83 | 4.93 | 0.573 | 1.04 | 1.09 | 0.14 |
| Kp4831 | 0.92 | 4.76 | 0.358 | 0.95 | 0.75 | 0.096 |
| Kp5022 | 0.56 | 2.82 | 0.696 | 1.81 | 0.39 | 0.056 |
| Kp5256 | 0.53 | 3.04 | 0.55 | 1.12 | 0.35 | 0.046 |
| Kp5712 | 0.56 | 3.98 | 0.65 | 1.61 | 0.49 | 0.071 |
| Kp6486 | 0.89 | 4.21 | 0.591 | 0.69 | 0.91 | 0.113 |
| Kp6918 | 0.99 | 3.72 | 0.618 | 0.79 | 1.03 | 0.153 |
| Kp7191 | 0.6 | 4.11 | 0.478 | 0.77 | 1.04 | 0.148 |
| Kp7637 | 0.54 | 4.02 | 0.608 | 0.82 | 1.33 | 0.177 |
| Kp7951 | 0.55 | 2.64 | 0.63 | 0.84 | 0.92 | 0.15 |
| Kp8039 | 0.58 | 4.59 | 0.723 | 0.67 | 1.78 | 0.231 |
| Kp8341 | 0.82 | 3.74 | 0.573 | 0.81 | 0.72 | 0.089 |
| Kp8715 | 0.6 | 2.81 | 0.358 | 1.06 | 0.26 | 0.035 |
| Kp10072 | 0.68 | 3.73 | 0.696 | 0.77 | 0.91 | 0.115 |
| Kp10079 | 0.49 | 2.6 | 0.55 | 0.9 | 0.98 | 0.144 |
| Kp10081 | 0.69 | 3.68 | 0.65 | 0.9 | 0.84 | 0.112 |
| Kp10403 | 0.88 | 3.23 | 0.591 | 1.05 | 0.68 | 0.097 |
| Kp10856 | 0.52 | 3.29 | 0.618 | 0.86 | 0.51 | 0.075 |
| Kp11465 | 0.79 | 2.59 | 0.478 | 1.17 | 0.57 | 0.084 |
| Kp11469 | 0.66 | 3.55 | 0.608 | 0.78 | 0.65 | 0.081 |
| Kp11997 | 0.7 | 3.6 | 0.63 | 1.63 | 0.97 | 0.107 |
| Kp12022 | 0.73 | 4.3 | 0.723 | 0.94 | 1.82 | 0.204 |

## REFERENCES

1. Klevens RM, Edwards JR, Richards CL, Jr., Horan TC, Gaynes RP, Pollock DA, Cardo DM. Estimating health care-associated infections and deaths in U.S. hospitals, 2002. Public Health Rep. 2007;122(2):160–6. doi: 10.1177/003335490712200205. PubMed PMID: 17357358; PMCID: PMC1820440.

2. Yang X, Chen H, Zheng Y, Qu S, Wang H, Yi F. Disease burden and long-term trends of urinary tract infections: A worldwide report. Front Public Health. 2022;10:888205. Epub 20220727. doi: 10.3389/fpubh.2022.888205. PubMed PMID: 35968451; PMCID: PMC9363895.

3. Caneiras C, Lito L, Melo-Cristino J, Duarte A. Community- and Hospital-Acquired Klebsiella pneumoniae Urinary Tract Infections in Portugal: Virulence and Antibiotic Resistance. Microorganisms. 2019;7(5). Epub 20190516. doi: 10.3390/microorganisms7050138. PubMed PMID: 31100810; PMCID: PMC6560439.

4. Frimodt-Moller N, Bjerrum L. Treating urinary tract infections in the era of antibiotic resistance. Expert Rev Anti Infect Ther. 2023;21(12):1301–8. Epub 20231124. doi: 10.1080/14787210.2023.2279104. PubMed PMID: 37922147.

5. Flores-Mireles AL, Walker JN, Caparon M, Hultgren SJ. Urinary tract infections: epidemiology, mechanisms of infection and treatment options. Nat Rev Microbiol. 2015;13(5):269–84. Epub 20150408. doi: 10.1038/nrmicro3432. PubMed PMID: 25853778; PMCID: PMC4457377.

6. (WHO bacterial priority pathogens list, 2024: Bacterial pathogens of public health importance to guide research, development and strategies to prevent and control antimicrobial resistance. World Health Organization, 2024 17 May 2024. Report No.: 978-92-4-009346-1.

7. Gorrie CL, Mirceta M, Wick RR, Judd LM, Lam MMC, Gomi R, Abbott IJ, Thomson NR, Strugnell RA, Pratt NF, Garlick JS, Watson KM, Hunter PC, Pilcher DV, McGloughlin SA, Spelman DW, Wyres KL, Jenney AWJ, Holt KE. Genomic dissection of Klebsiella pneumoniae infections in hospital patients reveals insights into an opportunistic pathogen. Nat Commun. 2022;13(1):3017. Epub 20220531. doi: 10.1038/s41467-022-30717-6. PubMed PMID: 35641522; PMCID: PMC9156735.

8. Podschun R, Ullmann U. Klebsiella spp. as nosocomial pathogens: epidemiology, taxonomy, typing methods, and pathogenicity factors. Clin Microbiol Rev. 1998;11(4):589–603. doi: 10.1128/CMR.11.4.589. PubMed PMID: 9767057; PMCID: PMC88898.

9. Shon AS, Bajwa RP, Russo TA. Hypervirulent (hypermucoviscous) Klebsiella pneumoniae: a new and dangerous breed. Virulence. 2013;4(2):107–18. Epub 20130109. doi: 10.4161/viru.22718. PubMed PMID: 23302790; PMCID: PMC3654609.

10. Marr CM, Russo TA. Hypervirulent Klebsiella pneumoniae: a new public health threat. Expert Rev Anti Infect Ther. 2019;17(2):71–3. Epub 20181205. doi: 10.1080/14787210.2019.1555470. PubMed PMID: 30501374; PMCID: PMC6349525.

11. Paczosa MK, Mecsas J. Klebsiella pneumoniae: Going on the Offense with a Strong Defense. Microbiol Mol Biol Rev. 2016;80(3):629–61. Epub 20160615. doi: 10.1128/MMBR.00078-15. PubMed PMID: 27307579; PMCID: PMC4981674.

12. Foxman B, Brown P. Epidemiology of urinary tract infections: transmission and risk factors, incidence, and costs. Infect Dis Clin North Am. 2003;17(2):227–41. doi: 10.1016/s0891-5520(03)00005-9. PubMed PMID: 12848468.

13. Kaper JB, Nataro JP, Mobley HL. Pathogenic Escherichia coli. Nat Rev Microbiol. 2004;2(2):123–40. doi: 10.1038/nrmicro818. PubMed PMID: 15040260.

14. Connell I, Agace W, Klemm P, Schembri M, Marild S, Svanborg C. Type 1 fimbrial expression enhances Escherichia coli virulence for the urinary tract. Proc Natl Acad Sci U S A. 1996;93(18):9827–32. doi: 10.1073/pnas.93.18.9827. PubMed PMID: 8790416; PMCID: PMC38514.

15. Stahlhut SG, Chattopadhyay S, Struve C, Weissman SJ, Aprikian P, Libby SJ, Fang FC, Krogfelt KA, Sokurenko EV. Population variability of the FimH type 1 fimbrial adhesin in Klebsiella pneumoniae. J Bacteriol. 2009;191(6):1941–50. Epub 20090116. doi: 10.1128/JB.00601-08. PubMed PMID: 19151141; PMCID: PMC2648365.

16. Lopatto EDB, Pinkner JS, Sanick DA, Potter RF, Liu LX, Bazan Villicana J, Tamadonfar KO, Ye Y, Zimmerman MI, Gualberto NC, Dodson KW, Janetka JW, Hunstad DA, Hultgren SJ. Conformational ensembles in Klebsiella pneumoniae FimH impact uropathogenesis. Proc Natl Acad Sci U S A. 2024;121(39):e2409655121. Epub 20240917. doi: 10.1073/pnas.2409655121. PubMed PMID: 39288182; PMCID: PMC11441496.

17. Nassif X, Fournier JM, Arondel J, Sansonetti PJ. Mucoid phenotype of Klebsiella pneumoniae is a plasmid-encoded virulence factor. Infect Immun. 1989;57(2):546–52. doi: 10.1128/iai.57.2.546-552.1989. PubMed PMID: 2643575; PMCID: PMC313131.

18. Bengoechea JA, Sa Pessoa J. Klebsiella pneumoniae infection biology: living to counteract host defences. FEMS Microbiol Rev. 2019;43(2):123–44. doi: 10.1093/femsre/fuy043. PubMed PMID: 30452654; PMCID: PMC6435446.

19. Tarkkanen AM, Allen BL, Williams PH, Kauppi M, Haahtela K, Siitonen A, Orskov I, Orskov F, Clegg S, Korhonen TK. Fimbriation, capsulation, and iron-scavenging systems of Klebsiella strains associated with human urinary tract infection. Infect Immun. 1992;60(3):1187–92. doi: 10.1128/iai.60.3.1187-1192.1992. PubMed PMID: 1347287; PMCID: PMC257611.

20. Ovchinnikova OG, Treat LP, Teelucksingh T, Clarke BR, Miner TA, Whitfield C, Walker KA, Miller VL. Hypermucoviscosity Regulator RmpD Interacts with Wzc and Controls Capsular Polysaccharide Chain Length. mBio. 2023;14(3):e0080023. Epub 20230504. doi: 10.1128/mbio.00800-23. PubMed PMID: 37140436; PMCID: PMC10294653.

21. Khadka S, Ring BE, Walker RS, Krzeminski LR, Pariseau DA, Hathaway M, Mobley HLT, Mike LA. Urine-mediated suppression of Klebsiella pneumoniae mucoidy is counteracted by spontaneous Wzc variants altering capsule chain length. mSphere. 2023;8(5):e0028823. Epub 20230823. doi: 10.1128/msphere.00288-23. PubMed PMID: 37610214; PMCID: PMC10597399.

22. Ernst CM, Braxton JR, Rodriguez-Osorio CA, Zagieboylo AP, Li L, Pironti A, Manson AL, Nair AV, Benson M, Cummins K, Clatworthy AE, Earl AM, Cosimi LA, Hung DT. Adaptive evolution of virulence and persistence in carbapenem-resistant Klebsiella pneumoniae. Nat Med. 2020;26(5):705–11. Epub 20200413. doi: 10.1038/s41591-020-0825-4. PubMed PMID: 32284589; PMCID: PMC9194776.

23. Martin RM, Cao J, Brisse S, Passet V, Wu W, Zhao L, Malani PN, Rao K, Bachman MA. Molecular Epidemiology of Colonizing and Infecting Isolates of Klebsiella pneumoniae. mSphere. 2016;1(5). Epub 20161019. doi: 10.1128/mSphere.00261-16. PubMed PMID: 27777984; PMCID: PMC5071533.

24. Vornhagen J, Roberts EK, Unverdorben L, Mason S, Patel A, Crawford R, Holmes CL, Sun Y, Teodorescu A, Snitkin ES, Zhao L, Simner PJ, Tamma PD, Rao K, Kaye KS, Bachman MA. Combined comparative genomics and clinical modeling reveals plasmid-encoded genes are independently associated with Klebsiella infection. Nat Commun. 2022;13(1):4459. Epub 20220801. doi: 10.1038/s41467-022-31990-1. PubMed PMID: 35915063; PMCID: PMC9343666.

25. Rao K, Patel A, Sun Y, Vornhagen J, Motyka J, Collingwood A, Teodorescu A, Baang JH, Zhao L, Kaye KS, Bachman MA. Risk Factors for Klebsiella Infections among Hospitalized Patients with Preexisting Colonization. mSphere. 2021;6(3):e0013221. Epub 20210623. doi: 10.1128/mSphere.00132-21. PubMed PMID: 34160237; PMCID: PMC8265626.

26. CDC H. Urinary Tract Infection (Catheter-Associated Urinary TractInfection [CAUTI] and Non-Catheter-Associated Urinary Tract Infection [UTI]) Events. CDC, 2026 Janaury 2026. Report No.

27. Wyres KL, Wick RR, Gorrie C, Jenney A, Follador R, Thomson NR, Holt KE. Identification of Klebsiella capsule synthesis loci from whole genome data. Microb Genom. 2016;2(12):e000102. Epub 20161212. doi: 10.1099/mgen.0.000102. PubMed PMID: 28348840; PMCID: PMC5359410.

28. Wick RR, Heinz E, Holt KE, Wyres KL. Kaptive Web: User-Friendly Capsule and Lipopolysaccharide Serotype Prediction for Klebsiella Genomes. J Clin Microbiol. 2018;56(6). Epub 20180525. doi: 10.1128/JCM.00197-18. PubMed PMID: 29618504; PMCID: PMC5971559.

29. Lam MMC, Wick RR, Judd LM, Holt KE, Wyres KL. Kaptive 2.0: updated capsule and lipopolysaccharide locus typing for the Klebsiella pneumoniae species complex. Microb Genom. 2022;8(3). doi: 10.1099/mgen.0.000800. PubMed PMID: 35311639; PMCID: PMC9176290.

30. Walker KA, Treat LP, Sepulveda VE, Miller VL. The Small Protein RmpD Drives Hypermucoviscosity in Klebsiella pneumoniae. mBio. 2020;11(5). Epub 20200922. doi: 10.1128/mBio.01750-20. PubMed PMID: 32963003; PMCID: PMC7512549.

31. Kanehisa M, Goto S. KEGG: kyoto encyclopedia of genes and genomes. Nucleic Acids Res. 2000;28(1):27–30. doi: 10.1093/nar/28.1.27. PubMed PMID: 10592173; PMCID: PMC102409.

32. Buffet A, Rocha EPC, Rendueles O. Nutrient conditions are primary drivers of bacterial capsule maintenance in Klebsiella. Proc Biol Sci. 2021;288(1946):20202876. Epub 20210303. doi: 10.1098/rspb.2020.2876. PubMed PMID: 33653142; PMCID: PMC7935059.

33. Khadka S, Ring BE, Pariseau DA, Mike LA. Characterization of Klebsiella pneumoniae Extracellular Polysaccharides. Curr Protoc. 2023;3(11):e937. doi: 10.1002/cpz1.937. PubMed PMID: 38010271; PMCID: PMC10683871.

34. Kinney EL, Stark DJ, Khadka S, Tin CM, Hand TW, Bain W, Mike LA. Connections between Klebsiella pneumoniae bloodstream dynamics and serotype-independent capsule properties. Infect Immun. 2026;94(3):e0064125. Epub 20260129. doi: 10.1128/iai.00641-25. PubMed PMID: 41609496; PMCID: PMC12974125.

35. Salisbury SM, Miner TA, Kent LA, Lam MMC, Holt KE, Miller VL, Walker KA. The acquisition of rmpADC can increase virulence of classical Klebsiella pneumoniae in the absence of other hypervirulence-associated genes. mBio. 2026;17(1):e0312225. Epub 20251209. doi: 10.1128/mbio.03122-25. PubMed PMID: 41363447; PMCID: PMC12802315.

36. Mike LA, Stark AJ, Forsyth VS, Vornhagen J, Smith SN, Bachman MA, Mobley HLT. A systematic analysis of hypermucoviscosity and capsule reveals distinct and overlapping genes that impact Klebsiella pneumoniae fitness. PLoS Pathog. 2021;17(3):e1009376. Epub 20210315. doi: 10.1371/journal.ppat.1009376. PubMed PMID: 33720976; PMCID: PMC7993769.

37. Ryan BE, Holmes CL, Stark DJ, Shepard GE, Mills EG, Khadka S, Van Tyne D, Bachman MA, Mike LA. Arginine regulates the mucoid phenotype of hypervirulent Klebsiella pneumoniae. Nat Commun. 2025;16(1):5875. Epub 20250701. doi: 10.1038/s41467-025-61047-y. PubMed PMID: 40595687; PMCID: PMC12219000.

38. Khadka S, Gates G, Stark DJ, Allison B, Mike LA. Sugar Import Suppresses Klebsiella pneumoniae Mucoidy in cAMP-CRP-dependent Manner. bioRxiv. 2025. Epub 20251104. doi: 10.1101/2025.11.04.686645. PubMed PMID: 41279589; PMCID: PMC12637589.

39. Wacharotayankun R, Arakawa Y, Ohta M, Tanaka K, Akashi T, Mori M, Kato N. Enhancement of extracapsular polysaccharide synthesis in Klebsiella pneumoniae by RmpA2, which shows homology to NtrC and FixJ. Infect Immun. 1993;61(8):3164–74. doi: 10.1128/iai.61.8.3164-3174.1993. PubMed PMID: 8335346; PMCID: PMC280984.

40. Le MN, Kayama S, Wyres KL, Yu L, Hisatsune J, Suzuki M, Yahara K, Terachi T, Sawa K, Takahashi S, Okuhara T, Kohama K, Holt KE, Mizutani T, Ohge H, Sugai M. Genomic epidemiology and temperature dependency of hypermucoviscous Klebsiella pneumoniae in Japan. Microb Genom. 2022;8(5). doi: 10.1099/mgen.0.000827. PubMed PMID: 35622495; PMCID: PMC9465067.

41. Russo TA, Olson R, Fang CT, Stoesser N, Miller M, MacDonald U, Hutson A, Barker JH, La Hoz RM, Johnson JR. Identification of Biomarkers for Differentiation of Hypervirulent Klebsiella pneumoniae from Classical K. pneumoniae. J Clin Microbiol. 2018;56(9). Epub 20180827. doi: 10.1128/JCM.00776-18. PubMed PMID: 29925642; PMCID: PMC6113484.

42. Maheswari UB, Palvai S, Anuradha PR, Kammili N. Hemagglutination and biofilm formation as virulence markers of uropathogenic Escherichia coli in acute urinary tract infections and urolithiasis. Indian J Urol. 2013;29(4):277–81. doi: 10.4103/0970-1591.120093. PubMed PMID: 24235787; PMCID: PMC3822341.

43. Shea AE, Frick-Cheng AE, Smith SN, Mobley HLT. Phenotypic Assessment of Clinical Escherichia coli Isolates as an Indicator for Uropathogenic Potential. mSystems. 2022;7(6):e0082722. Epub 20221129. doi: 10.1128/msystems.00827-22. PubMed PMID: 36445110; PMCID: PMC9765037.

44. McCabe WR, Kaijser B, Olling S, Uwaydah M, Hanson LA. Escherichia coli in bacteremia: K and O antigens and serum sensitivity of strains from adults and neonates. J Infect Dis. 1978;138(1):33–41. doi: 10.1093/infdis/138.1.33. PubMed PMID: 355575.

45. Coggon CF, Jiang A, Goh KGK, Henderson IR, Schembri MA, Wells TJ. A Novel Method of Serum Resistance by Escherichia coli That Causes Urosepsis. mBio. 2018;9(3). Epub 20180626. doi: 10.1128/mBio.00920-18. PubMed PMID: 29946047; PMCID: PMC6020292.

46. Doorduijn DJ, Rooijakkers SH, van Schaik W, Bardoel BW. Complement resistance mechanisms of Klebsiella pneumoniae. Immunobiology. 2016;221(10):1102–9. Epub 20160616. doi: 10.1016/j.imbio.2016.06.014. PubMed PMID: 27364766.

47. Short FL, Di Sario G, Reichmann NT, Kleanthous C, Parkhill J, Taylor PW. Genomic Profiling Reveals Distinct Routes To Complement Resistance in Klebsiella pneumoniae. Infect Immun. 2020;88(8). Epub 20200721. doi: 10.1128/IAI.00043-20. PubMed PMID: 32513855; PMCID: PMC7375759.

48. Martinez JJ, Mulvey MA, Schilling JD, Pinkner JS, Hultgren SJ. Type 1 pilus-mediated bacterial invasion of bladder epithelial cells. EMBO J. 2000;19(12):2803–12. doi: 10.1093/emboj/19.12.2803. PubMed PMID: 10856226; PMCID: PMC203355.

49. Dhakal BK, Mulvey MA. Uropathogenic Escherichia coli invades host cells via an HDAC6-modulated microtubule-dependent pathway. J Biol Chem. 2009;284(1):446–54. Epub 20081106. doi: 10.1074/jbc.M805010200. PubMed PMID: 18996840; PMCID: PMC2610520.

50. King JE, Aal Owaif HA, Jia J, Roberts IS. Phenotypic Heterogeneity in Expression of the K1 Polysaccharide Capsule of Uropathogenic Escherichia coli and Downregulation of the Capsule Genes during Growth in Urine. Infect Immun. 2015;83(7):2605–13. Epub 20150413. doi: 10.1128/IAI.00188-15. PubMed PMID: 25870229; PMCID: PMC4468546.

51. Liu X, Xu Q, Yang X, Heng H, Yang C, Yang G, Peng M, Chan E-C, Chen S. Capsular polysaccharide enables Klebsiella pneumoniae to evade phagocytosis by blocking host-bacteria interactions. mBio. 2025;16(3):e0383824. Epub 20250214. doi: 10.1128/mbio.03838-24. PubMed PMID: 39950808; PMCID: PMC11898582.

52. Connell H, Poulsen LK, Klemm P. Expression of type 1 and P fimbriae in situ and localisation of a uropathogenic Escherichia coli strain in the murine bladder and kidney. Int J Med Microbiol. 2000;290(7):587–97. doi: 10.1016/S1438-4221(00)80006-5. PubMed PMID: 11200540.

53. Arthur M, Johnson CE, Rubin RH, Arbeit RD, Campanelli C, Kim C, Steinbach S, Agarwal M, Wilkinson R, Goldstein R. Molecular epidemiology of adhesin and hemolysin virulence factors among uropathogenic Escherichia coli. Infect Immun. 1989;57(2):303–13. doi: 10.1128/iai.57.2.303-313.1989. PubMed PMID: 2563254; PMCID: PMC313098.

54. Schaeffer AJ, Schwan WR, Hultgren SJ, Duncan JL. Relationship of type 1 pilus expression in Escherichia coli to ascending urinary tract infections in mice. Infect Immun. 1987;55(2):373–80. doi: 10.1128/iai.55.2.373-380.1987. PubMed PMID: 2879794; PMCID: PMC260337.

55. Xia Y, Gally D, Forsman-Semb K, Uhlin BE. Regulatory cross-talk between adhesin operons in Escherichia coli: inhibition of type 1 fimbriae expression by the PapB protein. EMBO J. 2000;19(7):1450–7. doi: 10.1093/emboj/19.7.1450. PubMed PMID: 10747013; PMCID: PMC310214.

56. Lane MC, Mobley HL. Role of P-fimbrial-mediated adherence in pyelonephritis and persistence of uropathogenic Escherichia coli (UPEC) in the mammalian kidney. Kidney Int. 2007;72(1):19–25. Epub 20070328. doi: 10.1038/sj.ki.5002230. PubMed PMID: 17396114.

57. Gunther NWt, Lockatell V, Johnson DE, Mobley HL. In vivo dynamics of type 1 fimbria regulation in uropathogenic Escherichia coli during experimental urinary tract infection. Infect Immun. 2001;69(5):2838–46. doi: 10.1128/IAI.69.5.2838-2846.2001. PubMed PMID: 11292696; PMCID: PMC98232.

58. van der Bosch JF, Verboom-Sohmer U, Postma P, de Graaff J, MacLaren DM. Mannose-sensitive and mannose-resistant adherence to human uroepithelial cells and urinary virulence of Escherichia coli. Infect Immun. 1980;29(1):226–33. doi: 10.1128/iai.29.1.226-233.1980. PubMed PMID: 6105132; PMCID: PMC551100.

59. Gander RM, Thomas VL, Forland M. Mannose-resistant hemagglutination and P receptor recognition of uropathogenic Escherichia coli isolated from adult patients. J Infect Dis. 1985;151(3):508–13. doi: 10.1093/infdis/151.3.508. PubMed PMID: 2857751.

60. Rosen DA, Pinkner JS, Walker JN, Elam JS, Jones JM, Hultgren SJ. Molecular variations in Klebsiella pneumoniae and Escherichia coli FimH affect function and pathogenesis in the urinary tract. Infect Immun. 2008;76(7):3346–56. Epub 20080512. doi: 10.1128/IAI.00340-08. PubMed PMID: 18474655; PMCID: PMC2446687.

61. Hyun M, Lee JY, Kim HA, Ryu SY. Comparison of Escherichia coli and Klebsiella pneumoniae Acute Pyelonephritis in Korean Patients. Infect Chemother. 2019;51(2):130–41. doi: 10.3947/ic.2019.51.2.130. PubMed PMID: 31270992; PMCID: PMC6609746.

62. Pariseau DA, Ring BE, Khadka S, Mike LA. Cultivation and Genomic DNA Extraction of Klebsiella pneumoniae. Curr Protoc. 2024;4(1):e932. doi: 10.1002/cpz1.932. PubMed PMID: 38279957; PMCID: PMC11407547.

63. Argimon S, David S, Underwood A, Abrudan M, Wheeler NE, Kekre M, Abudahab K, Yeats CA, Goater R, Taylor B, Harste H, Muddyman D, Feil EJ, Brisse S, Holt K, Donado-Godoy P, Ravikumar KL, Okeke IN, Carlos C, Aanensen DM, Resistance NGHRUoGSoA. Rapid Genomic Characterization and Global Surveillance of Klebsiella Using Pathogenwatch. Clin Infect Dis. 2021;73(Suppl_4):S325–S35. doi: 10.1093/cid/ciab784. PubMed PMID: 34850838; PMCID: PMC8634497.

64. Lam MMC, Wick RR, Watts SC, Cerdeira LT, Wyres KL, Holt KE. A genomic surveillance framework and genotyping tool for Klebsiella pneumoniae and its related species complex. Nat Commun. 2021;12(1):4188. Epub 20210707. doi: 10.1038/s41467-021-24448-3. PubMed PMID: 34234121; PMCID: PMC8263825.

65. Ong CL, Beatson SA, Totsika M, Forestier C, McEwan AG, Schembri MA. Molecular analysis of type 3 fimbrial genes from Escherichia coli, Klebsiella and Citrobacter species. BMC Microbiol. 2010;10:183. Epub 20100624. doi: 10.1186/1471-2180-10-183. PubMed PMID: 20576143; PMCID: PMC2900259.

66. Olson RD, Assaf R, Brettin T, Conrad N, Cucinell C, Davis JJ, Dempsey DM, Dickerman A, Dietrich EM, Kenyon RW, Kuscuoglu M, Lefkowitz EJ, Lu J, Machi D, Macken C, Mao C, Niewiadomska A, Nguyen M, Olsen GJ, Overbeek JC, Parrello B, Parrello V, Porter JS, Pusch GD, Shukla M, Singh I, Stewart L, Tan G, Thomas C, VanOeffelen M, Vonstein V, Wallace ZS, Warren AS, Wattam AR, Xia F, Yoo H, Zhang Y, Zmasek CM, Scheuermann RH, Stevens RL. Introducing the Bacterial and Viral Bioinformatics Resource Center (BV-BRC): a resource combining PATRIC, IRD and ViPR. Nucleic Acids Res. 2023;51(D1):D678–D89. doi: 10.1093/nar/gkac1003. PubMed PMID: 36350631; PMCID: PMC9825582.

67. Anderson MT, Mitchell LA, Zhao L, Mobley HLT. Capsule Production and Glucose Metabolism Dictate Fitness during Serratia marcescens Bacteremia. mBio. 2017;8(3). Epub 20170523. doi: 10.1128/mBio.00740-17. PubMed PMID: 28536292; PMCID: PMC5442460.

68. Hagberg L, Engberg I, Freter R, Lam J, Olling S, Svanborg Eden C. Ascending, unobstructed urinary tract infection in mice caused by pyelonephritogenic Escherichia coli of human origin. Infect Immun. 1983;40(1):273–83. doi: 10.1128/iai.40.1.273-283.1983. PubMed PMID: 6339403; PMCID: PMC264845.

69. Hagberg L, Hull R, Hull S, Falkow S, Freter R, Svanborg Eden C. Contribution of adhesion to bacterial persistence in the mouse urinary tract. Infect Immun. 1983;40(1):265–72. doi: 10.1128/iai.40.1.265-272.1983. PubMed PMID: 6131870; PMCID: PMC264844.

70. Zychlinsky Scharff A, Albert ML, Ingersoll MA. Urinary Tract Infection in a Small Animal Model: Transurethral Catheterization of Male and Female Mice. J Vis Exp. 2017(130). Epub 20171201. doi: 10.3791/54432. PubMed PMID: 29286380; PMCID: PMC5755510.

